# Electrical Oscillations in Microtubules

**DOI:** 10.1101/2025.08.25.672199

**Authors:** Md Mohsin, Horacio F. Cantiello, María del Rocío Cantero, Marcelo Marucho

## Abstract

Environmental perturbations and local changes in cellular electric potential can stimulate cytoskeletal filaments to transmit ionic currents along their surface. Advanced models and accurate experiments may provide a molecular understanding of these processes and reveal their role in cell electrical activities. This article introduces a multi-scale electrokinetic model incorporating atomistic protein details and biological environments to characterize electrical impulses along microtubules. We consider that condensed ionic layers on microtubule surfaces form two coupled asymmetric nonlinear electrical transmission lines. The model accounts for tubulin-tubulin interactions, dissipation, and a nanopore coupling between inner and outer surfaces, enabling luminal currents, energy transfer, amplification, and oscillatory dynamics that resemble the experimentally observed transistor properties of microtubules. The approach has been used to analyze how different electrolyte conditions and voltage stimuli affect electrical impulses’ shape, attenuation, oscillation, and propagation velocity along microtubules. Integrating transistor-like properties in the microtubules model has profound implications for intracellular communication and bioelectronic applications.

## Introduction

Microtubules (MTs) are hollow cylindrical cytoskeleton polymers inside cells [1]. Apart from their well-known roles in controlling cell division, intracellular transport, and shape of the cells, MTs are dynamic biopolymers with remarkable electrical properties (Fig. 1).

**Figure 1.**
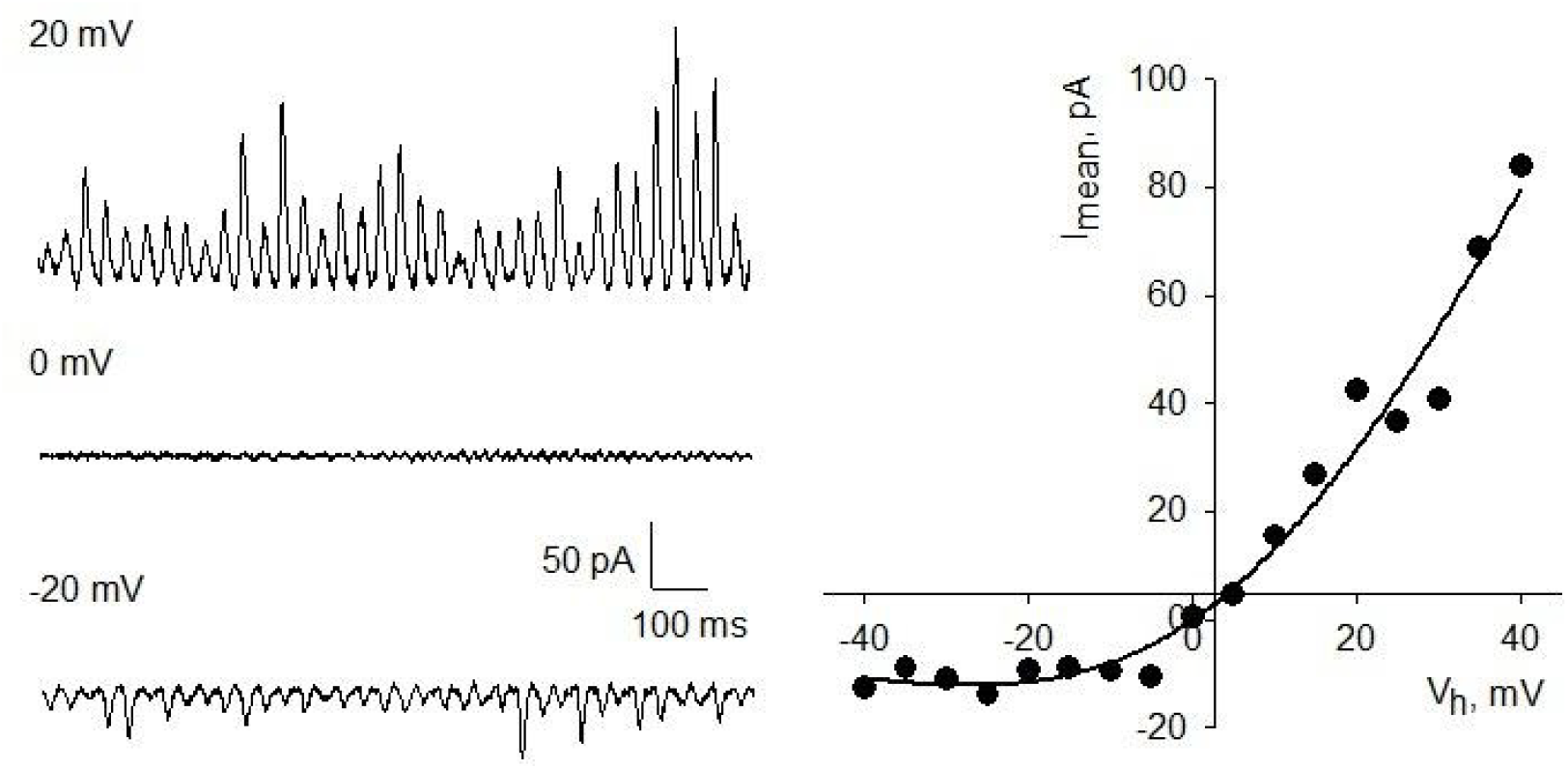
Electrical oscillations and rectification obtained in a two-dimensional MT structure, as described in reference [2]. (Left) Representative tracings of the effect of the input voltage, as indicated, on the MT sheet current oscillations. (Right) Current-to-voltage relationship obtained from integrated, mean currents of individual experiments without any changes in electrochemical gradient.

Recent studies from our group and others have demonstrated that MTs can behave as active bio-electronic elements rather than passive cytoskeletal scaffolds. Using patch-clamp electrophysiology, we reported for the first time that two-dimensional sheets of MTs exhibit spontaneous oscillations in ionic currents across a wide range of holding potentials, with robust spectral peaks at ∼39 Hz and higher-order modes [2]. These oscillations persisted across multiple preparations, suggesting that MT assemblies act as self-sustained oscillators. Subsequent work extended these findings to MT bundles, where coherent oscillations were observed in ensembles of brain-derived MTs, again with dominant frequencies in the gamma range (30–100 Hz) [3, 4]. More recently, isolated MTs stabilized with paclitaxel were shown to retain oscillatory behavior, confirming that the phenomenon is intrinsic to the MT polymer itself [5].

These oscillations are not merely biophysical curiosities but may be central to intracellular information processing. For example, electrical oscillations are observed in permeabilized neurites of cultured adult hippocampal neurons [4]. Unlike simple diffusion-based signaling, oscillatory ionic currents can provide frequency-encoded information, enabling MTs to function as biological signal processors. Oscillations at ∼39 Hz overlap with the gamma frequency band of brain electrical activity, raising the possibility that MTs contribute to higher-order neuronal signaling, synchronization, and cognition [6]. More broadly, intracellular oscillators provide a mechanism for amplifying weak perturbations, coordinating responses over long distances, and integrating multiple inputs, all of which are fundamental for cell signaling.

From a mechanistic perspective, we have proposed that MT oscillations emerge from the voltage-dependent gating properties of nanopores located at the interfaces between tubulin subunits. The wall of an MT is typically organized in what is known as a B-lattice structure, which features two distinct types of nanopores. Two additional nanopore types are also present in the alternative A-lattice structure [7]. Nanopore type 1 is located where the interdimer α/β interface of one tubulin dimer is adjacent to the interdimer α/β interface of a neighboring dimer. Nanopore type 2, in contrast, occurs where the intradimer β/α interface of one tubulin dimer lies next to the intradimer β/α interface of the adjacent dimer (Fig. 2). These pores are tapered, ∼4 nm in length, with diameters ranging from ∼0.7 to 1 nm [8], and are permeable primarily to cations [7, 9]. Although the biological roles of the nanopores and the MT lumen remain poorly understood, there is compelling evidence suggesting a functional relevance. For instance, taxane-based chemotherapeutics are known to diffuse through the MT wall via these nanopores to reach their luminal binding sites [7], and compounds such as cyclostreptin directly bind within nanopores [10], indicating that they are active conduits for ionic and molecular exchange. Moreover, recent findings that Gd^3+^ ions inhibit MT oscillations [11] further support the view that these nanopores behave as functional ionic gates that dynamically couple the luminal and surface electrochemical environments.

**Figure 2.**
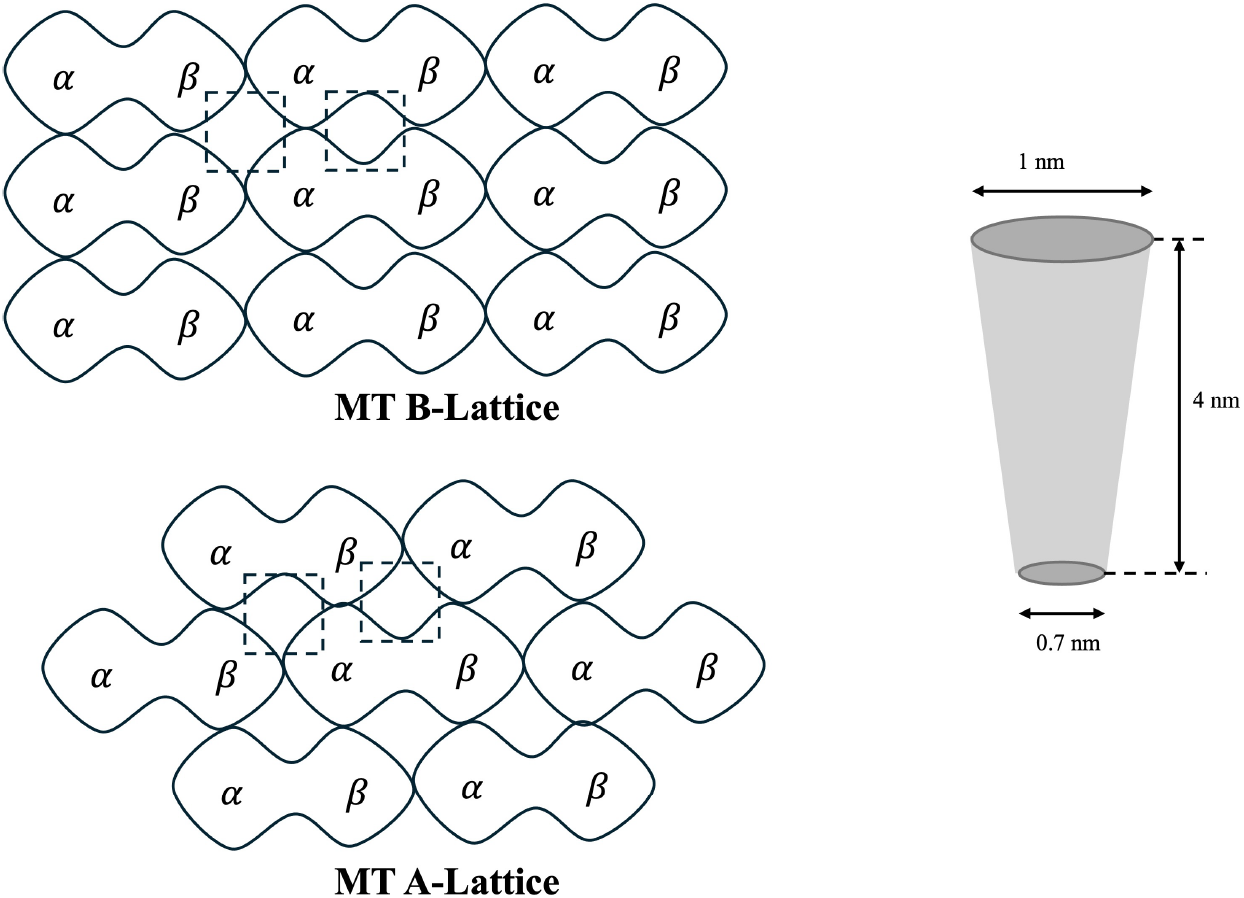
Schematic representation of microtubule lattice interfaces. (Left, top) B-lattice arrangement; (Left, bottom) A-lattice arrangement. Dashed boxes highlight the interdimer gaps corresponding to nanopores. These pores are illustrated by the funnel-shaped cartoon (right), representing possible ionic pathways across the microtubule wall.

Substantial work has been done to provide a molecular understanding of these experimental data on MTs. According to Manning’s theory [12, 13, 14], positive counterions condensate on their surface–a characteristic shared with other biomolecular filaments like DNA and F-actin–and move in response to applied electric fields in the form of ionic currents.

Pioneering approaches have modeled these biological electrical wires as single electrical transmission lines [8, 9, 15, 16], yielding non-perturbative Korteweg–de Vries (KdV) equations that predict ionic current propagating axially at near-uniform velocity. A more recent perturbative KdV model captured dissipation and damping contributions, predicting localized ionic wave packages (solitons) propagating with attenuated amplitudes and deceleration along actin filaments [17]. The approach also revealed that higher voltage inputs yield faster, narrower solitons, whereas lower inputs produce broader impulses. Meanwhile, intracellular environments exhibit lower electrical conductivity, higher capacitance, and broader double layers compared to the results obtained in in-vivo conditions. This multi-scale model also captured the impact of molecular conformational changes and solution protonation states, providing insights into pathological cytoskeletal conditions [18].

However, existing single transmission line models for MTs [8, 9, 15, 16, 19, 20] have overlooked the ionic current through the lumen, electrical signal amplification, and oscillations. Although the lumen side of the MT has a lower surface charge compared to the outer and, therefore, less condensed ions, the asymmetric conduction of the nanopores [7] and the tail oscillation [21] can produce a notable luminal ionic current comparable to the outer one. This current can play a key role in the electrical energy transfer between the inner and outer surfaces, leading to self-sustaining and self-reinforcing oscillating coupled solitons traveling along each surface. This phenomenon already manifests in two coupled nonlinear transmission lines (NLTLs) characterizing electrical networks in other fields [22, 23]. In these systems, solitons along each transmission line form bound state pairs, known as leapfrogging solitons (Fig. 3), propagate with opposite phases at nearly equal velocities, transfer electrical energy alternately from the leading soliton to the trailing soliton, giving rise to oscillations as they continuously overtake each other periodically.

**Figure 3.**
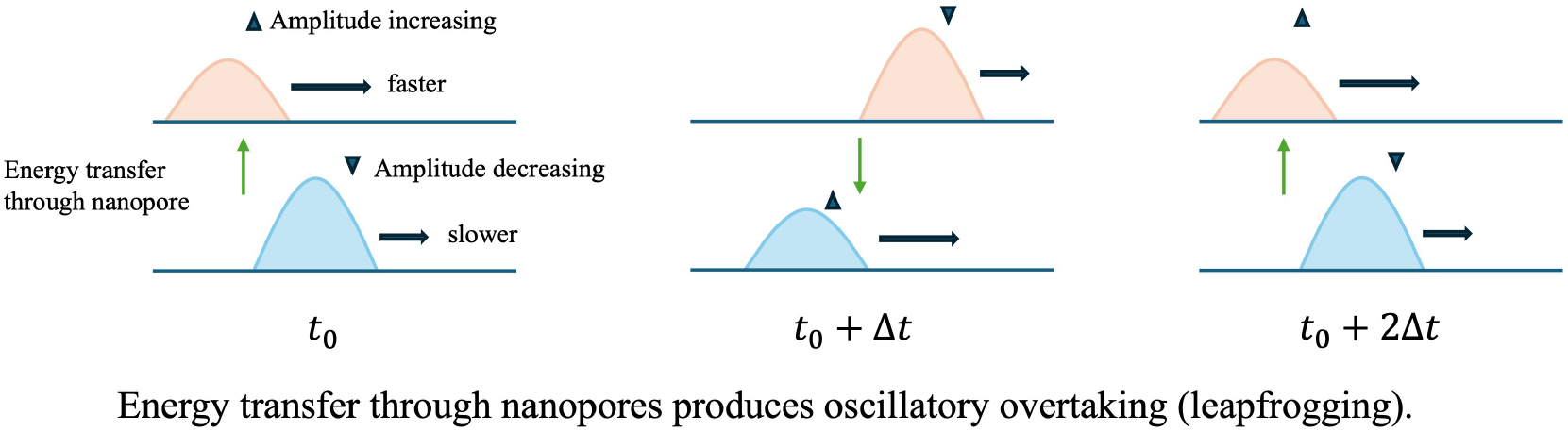
Schematic of leapfrogging soliton dynamics. Nanopore-mediated energy transfer produces oscillatory overtaking (‘leapfrogging’). When the outer soliton peak increases, the inner peak decreases, and vice versa. The growing soliton also propagates faster than the attenuating one, leading to oscillations and crossover events.

Moreover, in a transistor model for MT oscillations, the concept of time-variant pore conductance aligns naturally with electrochemical gating, where the nanopores function as dynamic control elements, similar to the gate of a transistor.

In our earlier studies, patch-clamp recordings of MTs revealed voltage-dependent amplification [24] and state-dependent conductance with hysteresis [25], two hallmark behaviors of transistor and memristive devices, respectively. These findings suggest that MT nanopores may function as dynamic gating elements, modulating ionic and electronic transport in a manner analogous to semiconductor components. Thus, the transistor/memristor analogies are mechanistically supported by nanopore gating and voltage-dependent conduction, rather than being purely conceptual.

In a Bipolar Junction Transistor (BJT) analogy (originally postulated in [24]), the base current would control the much larger collector-emitter current. In MTs, the nanopores’ conductance and small modulations in local ionic gating, such as protonation-deprotonation cycles or divalent cation binding, would lead to significant changes in transmembrane ionic flow, where the nonlinear gain observed in MTs resembles transistor-like amplification.

In a MOSFET, the gate voltage modulates the channel conductivity by controlling charge carrier density. In this case, the nanopores in the MT, acting as gates, could modulate ionic currents through local changes in charge distribution, possibly driven by binding/unbinding of ions or conformational shifts. In MTs, local ionic depletion or enrichment in the nanopores could control the current flow, acting as a dynamic gating mechanism, where cyclic shifts in depletion/enrichment regions, driven by feedback from the intrinsic electrical activity, could help emerge the observed oscillations. Additionally, MTs exhibit voltage-dependent resonance frequencies, suggesting an intrinsic field-effect behavior where the conductance oscillates due to internal charge-wave interactions.

Thus, the time-variant conductance of nanopores in MTs fits well into a transistor-based framework, particularly one where ionic gating modulates electrical properties dynamically. This supports the core idea that MTs function as biological semiconductors, where oscillatory gating enables information processing and amplification.

Based on these premises, in this article, we introduce a novel approach that considers condensed ionic layers on the outer and inner surfaces of the MT to form a pair of coupled nonlinear electrical transmission lines generating leapfrogging solitons. Additionally, counterions are exchanged between the inner and outer transmission lines through nanopores characterized by a nonlinear voltage-dependent resistance that resembles the transistor properties of MTs. The multi-scale electrokinetic model accounts for the atomistic details of the protein molecular structure and biological environment. The formulation includes non-trivial contributions to the ionic electrical conductivity and capacitance from the diffuse part of the tubulin’s ionic layers. It also accounts for the asymmetric outer and inner surface charge densities, the hollow size, a low dielectric constant in the condensed layers generated by the high polarization of immobile water molecules, and a lower dielectric constant between the outer and inner surfaces coming from the internal MT molecular structure properties. We utilized this tubulin characterization in an outer and inner nonlinear inhomogeneous transmission line prototype model to account for the tubulin–tubulin interactions, dissipation, and damping perturbations along the filament length. We used Kirchoff’s law to solve these electric circuits, obtaining two coupled perturbative KdV equations, one for the outer and one for the inner NLTL. We applied an adiabatic perturbation theory to transform these two third-order coupled differential equations [26] into four coupled first-order differential equations, two for the soliton phases and two for their amplitudes. We solved these equations numerically using Mathematica software. We also derived simple, easy-to-evaluate approximate analytical solutions for practical applications. Illustrative images and more details on the approach and the solution are provided in the Methods section and supplementary document. The approach has been used to determine the effects of electrolyte conditions and voltage stimulus on the electrical impulse shape, attenuation, oscillation, and propagation velocity along MTs.

## Results

In Fig. 4, we present the time evolution of the outer and inner (lumen) voltage signal solutions *V* (*x, t*) and *W* (*x, t*) for input voltages of 0.04 V and 0.08 V. The results display oscillatory traveling solitons at increasing (snapshot) distances *x* = 0.1, 2.6, 4.6, 6.6, 8.6, 10.6, 12.6, and 14.6 *µm* along the MT. The nanopores and the electrical double layer (EDL) on MT’s outer and lumen surfaces were characterized using the voltage-dependent nonlinear resistance *R*_*p*_(*V*), and the asymmetric condensed ionic layer parameters provided in the Methods section. They yield the following set of coupling parameter values: *M*_1_ = −2.450, *M*_2_ = 1.331, *M*_3_ = −1.060, *M*_4_ = 2.081, *N*_1_ = 0.0013, *N*_2_ = 0.0014, *N*_3_ = 0.0024, *N*_4_ = 0.0005, *F*_1_ = −0.102, *F*_2_ = −0.044; and *M*_1_ = −1.721, *M*_2_ = 0.912, *M*_3_ = −0.670, *M*_4_ = 1.402, *N*_1_ = 0.0011, *N*_2_ = 0.0009, *N*_3_ = 0.0021, *N*_4_ = 0.0003, *F*_1_ = −0.072, *F*_2_ = −0.028 for 0.04 V and 0.08 V input voltages respectively. These parameters appear in the main coupled soliton differential equations 12 and 13, and they are linked by equations S33–S44 in the supplementary document to the MT model parameter values. In the formulation, *M*_*i*_ denote the nanopore-mediated conductance and energy exchange between the outer and inner conduction pathways; *N*_*i*_ represent the dissipative losses linked to ionic conduction; and *F*_*i*_ correspond to higher-order dispersion.

**Figure 4.**
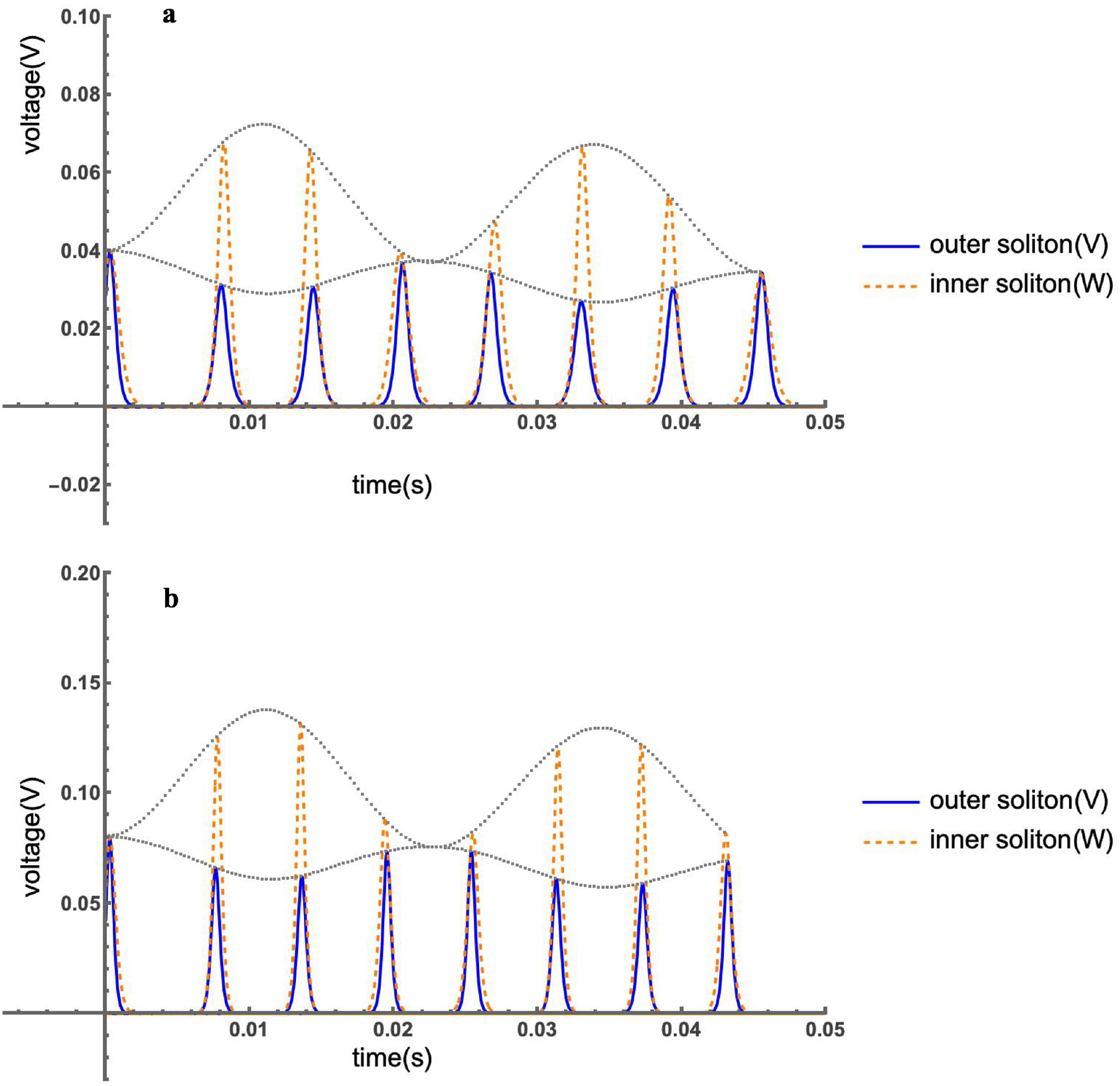
Soliton-like voltage propagation along the MT under asymmetric ionic conditions.Time evolution of the outer voltage *V* (*x, t*) and inner (lumen) voltage *W* (*x, t*) for input voltages of 0.04 V (a) and 0.08 V (b) at increasing positions along the microtubule: x = 0.1, 2.6, 4.6, 6.6, 8.6, 10.6, 12.6, 14.6 μm. Snapshot positions are chosen for visualization of equidistant soliton profiles. The black dotted lines are introduced to help visualize the time oscillation of the soliton amplitude peaks.

The soliton dynamics along the MT were obtained using Eq. 13 in Ref. [17] and the model parameter values mentioned above. The results are presented in Fig. 5, showing opposite-in-phase oscillations between the outer and inner soliton propagation velocities.

**Figure 5.**
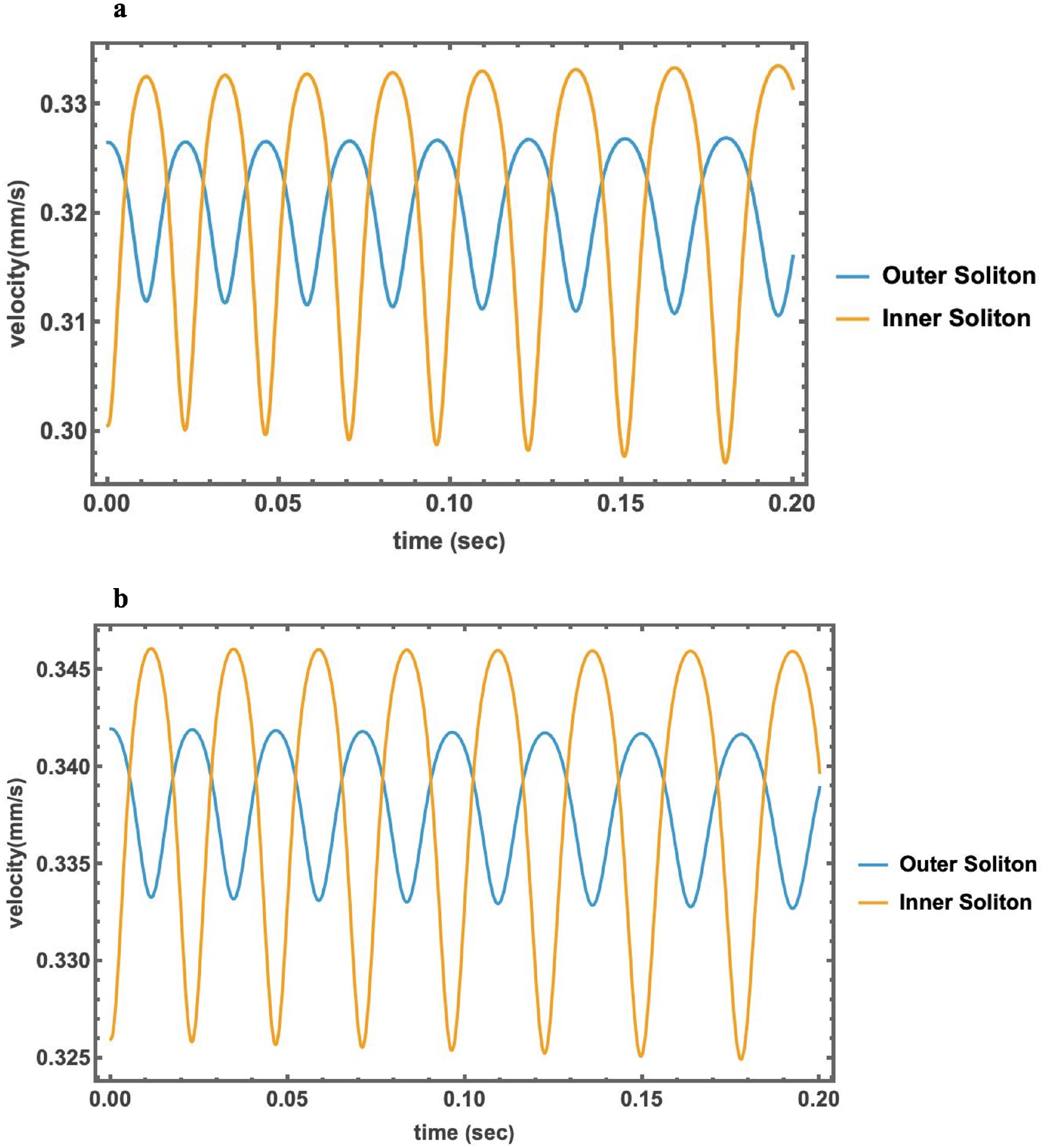
Soliton propagation velocities along the MT. (a) Input voltage 0.04 V. (b) Input voltage 0.08 V. These results correspond to the soliton-like voltage propagation described in Fig. 4.

Additionally, we determined the soliton oscillation frequency by analyzing the time-dependent signal derived from the numerical solutions of the MT model equations. We computed its numerical derivative to suppress DC and baseline drift, performed a discrete Fourier transform, and identified the frequency corresponding to the maximal amplitude of the one-sided spectrum. The dominant oscillation frequency was 39.1 Hz for both 0.04 V and 0.08 V input voltages, consistent with measurements from voltage-clamp experiments (Fig. 6).

**Figure 6.**
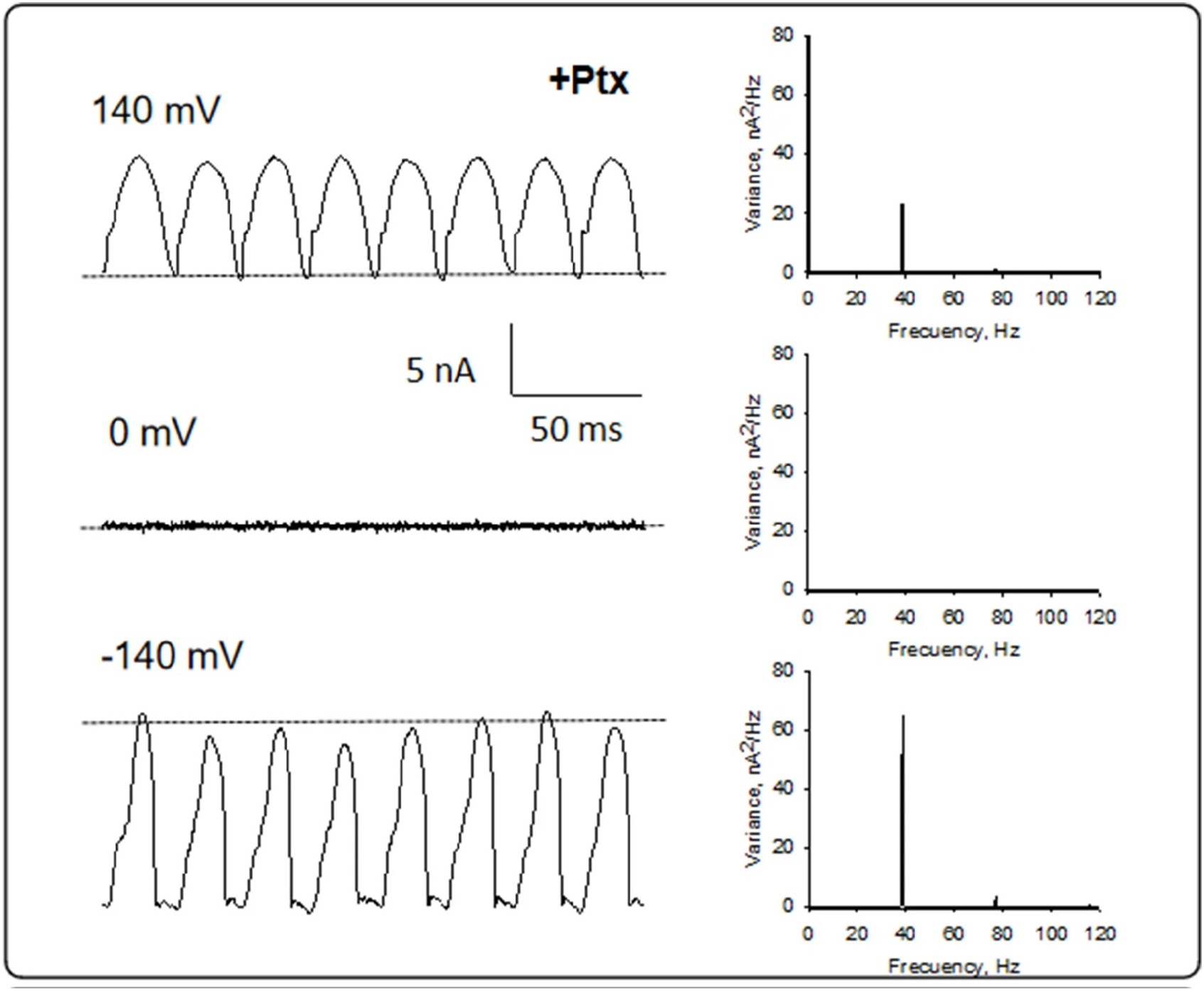
Experimental soliton oscillation frequencies. Current oscillations of Paclitaxel-stabilized isolated MTs. (Left) Current fluctuations at different holding potentials. (Right) Linear–linear plots of the Fourier power spectra obtained from tracings shown on the Left. The fundamental frequencies can be observed on the spectra (Ref. [5]).

To assess robustness, we systematically varied the nanopore resistance by ±10%, which shifted the oscillation frequency between 35.2–43.0 Hz for both 0.04 V and 0.08 V input voltages. Furthermore, a nonparametric bootstrap analysis (±10% random perturbation per data point, *n* = 300 runs) yielded 95% confidence interval of 35.2–41.0 Hz for 0.04 V and 35.2–43.0 Hz for 0.08 V. These analyses confirm that the predicted oscillatory behavior remains stable across plausible variations in pore resistance and input voltage.

The theoretical prediction obtained from the approximate analytical expression

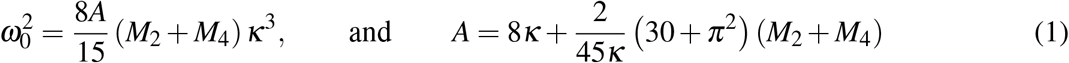

corresponds to a frequency of 43.3 Hz for 0.04 V and 41.9 Hz for 0.08 V. These values show close agreement with the numerical results while offering straightforward evaluation. The prediction at 0.08 V lies within the 95 % confidence interval of 35.2–43.0 Hz, while the 0.04 V estimate slightly exceeds its corresponding interval (35.2–41.0 Hz). These small deviations are consistent with the simplifying assumptions in the analytical derivation, which neglect higher-order nonlinear coupling terms captured in the full numerical solution. The parametes *M*_2_ and *M*_4_ in Eq. 1 are inversely proportional to the nanopore resistance *R*_*p*_. They are particularly crucial, as they reflect nanopore-mediated energy exchange between the inner and outer pathways, while *κ* defines the characteristic spatial scale of the soliton. Their contributions dominate the oscillation frequency and therefore the functional states of the nanopores. Pharmacological inhibition of nanopores corresponds mathematically to lowering *M*_2_ and *M*_4_, which leads to suppressed oscillations.

Moreover, we characterized the attenuation of the soliton amplitudes. We calculated the dimensionless damping parameter *γ*_0_ (see equations 18-22) using an exponential fitting curve 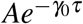 (where A is the amplitude and *τ* is the dimensionless time) and the aforementioned model parameter values. This damping parameter is proportional to the parameters *N*_*i*_ reflecting dissipative losses in the ionic double layer. The fitting results are presented in Fig. 7. The location of the soliton peaks are displayed in red points as a function of the dimensionless time *τ*, obtaining a value of *γ*_0_ = 1.23 × 10^−3^ for 0.04 V and *γ*_0_ = 1.19 × 10^−3^ for 0.08 V. These low damping values mean the solitons take ∼0.84 seconds to reduce their amplitudes by ∼37%. Robustness analyses showed that uniformly scaling the pore resistance by ±10% shifted the damping coefficient between (1.228–1.230) × 10^−3^ at 0.04 V and (1.171–1.663) × 10^−3^ at 0.08 V. In addition, a nonparametric bootstrap analysis (±10% random variation per data point, *n* = 300 runs) yielded a 95% confidence interval of (1.228–1.883) × 10^−3^ at 0.04 V and (1.172–1.702) × 10^−3^ at V, confirming that the attenuation dynamics remain robust to parameter variation.

**Figure 7.**
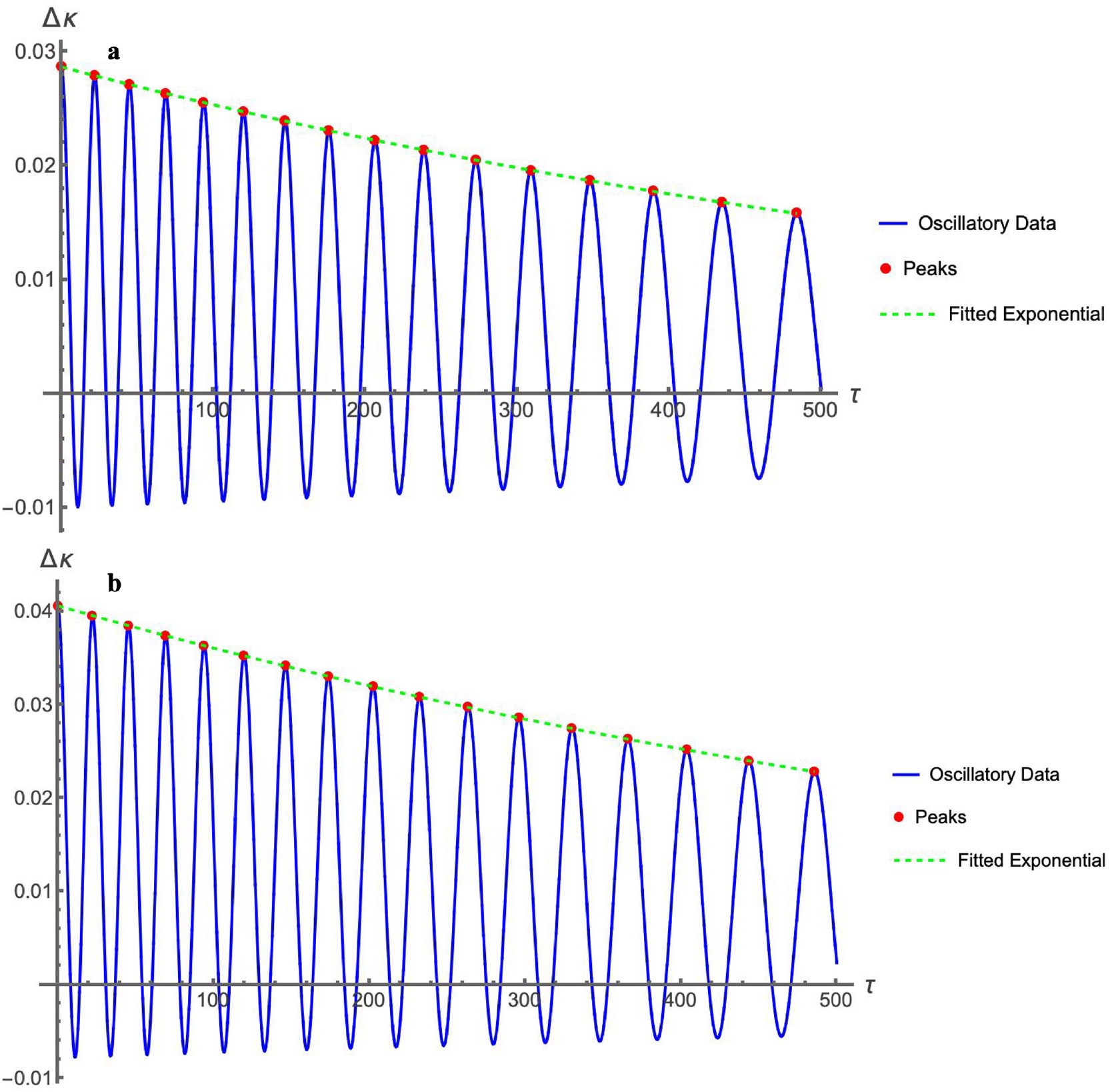
Soliton amplitude attenuation and damping parameter estimation. Exponential fitting of soliton peak amplitudes as a function of dimensionless time *τ*. (a) Input voltage 0.04 V. (b) Input voltage 0.08 V. Red points represent the soliton peak locations. These results correspond to the soliton-like voltage propagation described in Fig. 4.

In addition to uncertainty quantifications presented above, Figs. 8 and 9 display the sensitivity of the soliton dynamics to variations in the *M, N* and *F* parameters, demonstrating that the qualitative behavior is preserved across a broad parameter range.We examined the effects of the coupling and resistivity terms on the soliton oscillation frequency and amplitude by changing the model parameter values used previously to calculate the voltage signal solutions *V* (*x, t*) and *W* (*x, t*). In Fig. 8-a, we display the results obtained for the MT model parameter values *M*_1_ = 0, *M*_2_ = 1.331, *M*_3_ = 0, *M*_4_ = 2.081, *N*_1_ = 0.0013, *N*_2_ = 0, *N*_3_ = 0.0024, *N*_4_ = 0, *F*_1_ = 0, and *F*_2_ = 0; whereas, in Fig. 8-b, we show the results obtained for *M*_1_ = 0, *M*_2_ = 0.912, *M*_3_ = 0, *M*_4_ = 1.402, *N*_1_ = 0.0011, *N*_2_ = 0, *N*_3_ = 0.0021, *N*_4_ = 0, *F*_1_ = 0, and *F*_2_ = 0 (see equations 12 and 13 for more details on these parameters). Comparison between the results presented in Fig. 4 and 8 reveals a dominant role of the coupling terms *M*_2_ and *M*_4_ appearing in the soliton KdV equations 10 and 11 on the soliton oscillation. This explains why pharmacological blockade of nanopores by Taxol or Gd^3+^ suppresses oscillatory activity, reducing nanopore conductance lowers (*M*_2_ + *M*_4_), thereby eliminating the sustained oscillatory regime. Additionally, the soliton amplitudes are higher in Fig. 8 than in Fig. 4 because we have set some ionic resistivity terms in the soliton KdV equations equal to zero.

**Figure 8.**
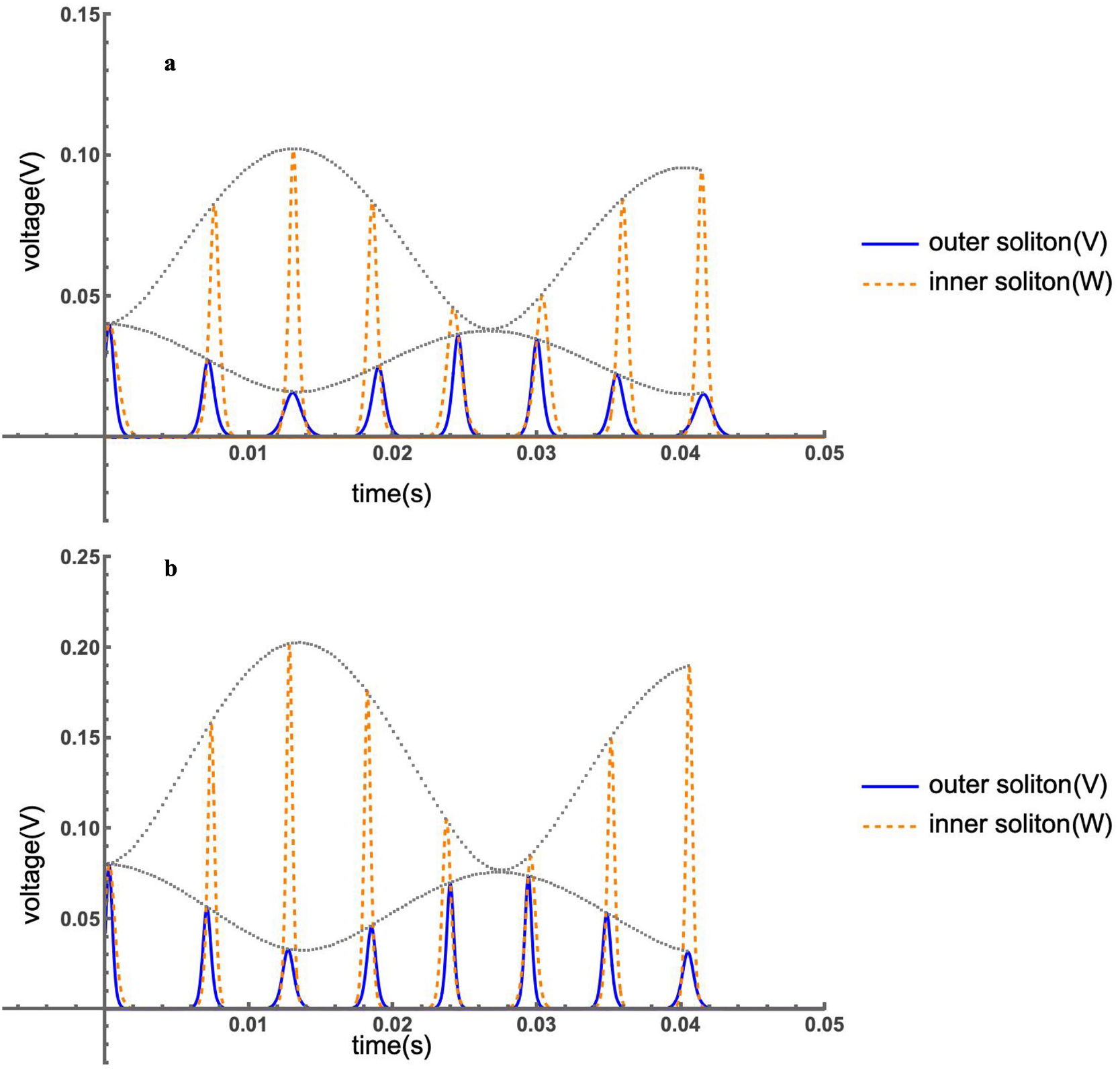
Effects of asymmetric ionic layer parameters on soliton propagation along MTs. (a) Input voltage 0.04 V. (b) Input voltage 0.08 V. Voltage signal solutions *V* (*x, t*) (outer) and *W* (*x, t*) (inner) are shown for asymmetric condensed layer parameter values different to those in Fig. 4. The black dotted line indicates the soliton amplitude peak oscillation.

**Figure 9.**
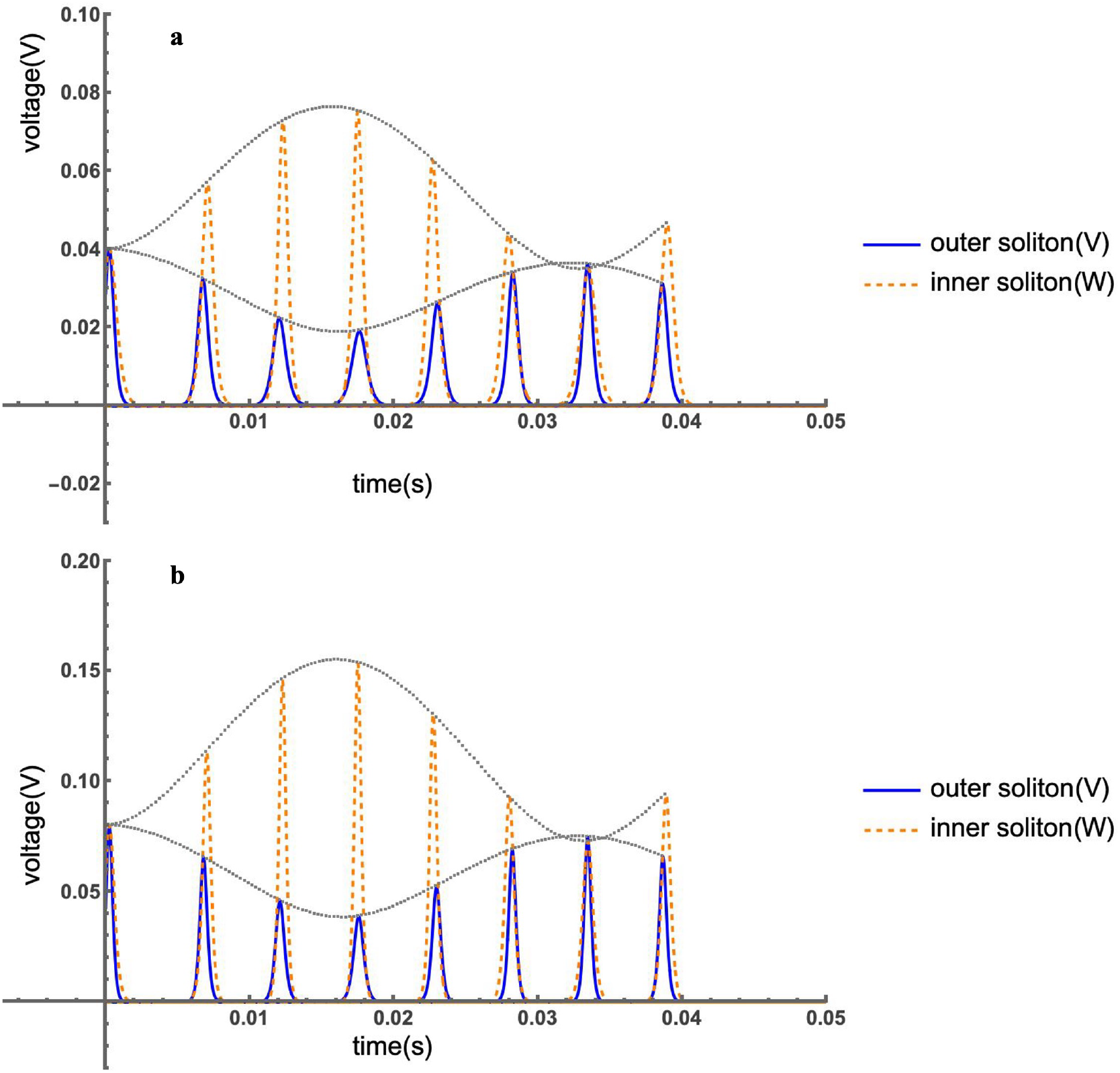
Effects of symmetric ionic layer parameters on soliton propagation along MTs. (a) Input voltage 0.04 V. (b) Input voltage 0.08 V. Voltage signal solutions *V* (*x, t*) (outer) and *W* (*x, t*) (inner) are shown for symmetric condensed ionic layer parameters. The black dotted line marks the soliton amplitude peak oscillation.

Finally, we elucidated the role of the asymmetry between the outer and inner condensed ionic layer parameters on the time evolution of the voltage signal solutions. Figures 9-a and 9-b display the results for the set of symmetric parameter values: *M*_1_ = *M*_2_ = *M*_3_ = *M*_4_ = 1.331, *N*_1_ = *N*_2_ = *N*_3_ = *N*_4_ = 0.0014, *F*_1_ = *F*_2_ = −0.044; and *M*_1_ = *M*_2_ = *M*_3_ = *M*_4_ = 0.912, *N*_1_ = *N*_2_ = *N*_3_ = *N*_4_ = 0.0009, *F*_1_ = *F*_2_ = −0.028, respectively. Comparison between the results shown in Fig. 9 and 4 demonstrates a substantial impact of the condensed ionic layer model parameter values on the soliton oscillation frequencies, velocities, and amplitudes.

In summary, our numerical results predict oscillatory electrical signal impulses propagation along the MT in the form of localized ionic wave packages, called leapfrogging solitons, for the range of voltage stimulus and electrolyte solutions typically present in in-vitro experimental conditions [2, 3, 5] (Fig. 4-7). While the outer soliton peak increases, the inner decreases, and vice versa (Fig. 4). Likewise, the velocity of the increasing soliton increases, whereas the velocity of the attenuating soliton decreases, and vice versa. Additionally, the outer and inner soliton velocities oscillate in opposite phases such that when one soliton accelerates, the other decelerates (Fig. 5). This oscillation frequency reproduces experimental measurements (Fig. 6). The electrical energy transferred from the leading soliton to the trailing soliton through the nanopores accelerates the trailing soliton by gaining kinetic energy. It decelerates the leading soliton by reducing kinetic energy. As a result, they shift their position at every oscillation cycle so that the leading one becomes the trailing soliton and vice versa. At lower input voltage, the average velocity and amplitude are lower; however, the velocity oscillation range is higher. Conversely, the average velocity and amplitude are higher at higher input voltage. Additionally, the velocity and amplitude changes are more pronounced in the inner soliton than in the outer one. This leapfrogging electrokinetic motion allows MTs to transmit electrical signals to relatively large distances with minimum power loss [22].

Further examination of the electrical signal propagation along MTs revealed a dominant role of the coupling terms between the outer and inner solitons on the oscillation frequency. In contrast, the diffusion and damping terms mainly affected the soliton amplitudes (Fig. 8). We noticed that the asymmetric characterization considered between the outer and inner condensed ionic layers of the MT plays a key role in the soliton solutions (Fig. 4 and 9). It generated more prominent outer and inner soliton amplitude changes and a slightly higher oscillation frequency than those obtained from a symmetric condensed ionic layer model.

Taken together, the coupling coefficients governing the model equations provide a quantitative handle on nanopore function. This theoretical framework is directly supported by experimental inhibition of nanopore conduction by Taxol and Gd^3+^, establishing a causal link between nanopore gating and sustained oscillatory behavior.

## Discussion

The present study provides a unified framework for understanding how MTs sustain and regulate electrical oscillations through their intrinsic nanostructure. Central to this behavior is the presence of nanopores located at the interfaces between α- and β-tubulin subunits. These nanopores form aqueous, ion-permeable channels that couple the lumen and the outer surface of the MT wall. Their voltage-dependent gating behavior introduces nonlinear conductance into the MT lattice, which is essential for the generation and propagation of oscillatory ionic currents.

### Nanopores as the structural origin of oscillatory activity

Previous work from our group has demonstrated that two-dimensional MT sheets exhibit spontaneous electrical oscillations under voltage-clamp conditions, even in the absence of external energy sources. These oscillations are frequency-rich, stable, and display entrainment under external stimuli, consistent with the presence of intrinsic feedback mechanisms within the polymer. The data strongly support the view that ionic exchange through nanopores— rather than through the open MT ends—is the key determinant of this activity. In this model, the nanopores act as ionic gates, alternating between conductive and non-conductive states in response to local electrostatic and mechanical conditions. The collective gating of these nanopores produces oscillatory currents that propagate along the MT, much like self-sustained waves in coupled transmission lines.

### Pharmacological evidence: inhibition by Taxol

The functional relevance of nanopores was further confirmed by the inhibitory effect of paclitaxel (Taxol) on MT electrical oscillations. Taxol binds to β-tubulin within the MT wall and stabilizes the polymer by suppressing dynamic instability. At the same time, its diffusion through the nanopores is essential for reaching its binding site, making these structures direct pharmacological targets. In our experiments, increasing concentrations of Taxol produced a dose-dependent reduction of oscillation amplitude, with complete inhibition at micromolar levels and a dissociation constant (K_D_) of ∼1.3 µM. Notably, this inhibition was voltage-dependent—raising the holding potential restored oscillatory behavior—demonstrating that the mechanism involves pore-blocking rather than polymer depolymerization. These findings confirm that ionic currents through nanopores are necessary for the oscillatory process and that interference with nanopore conduction abolishes the emergent electrical behavior of MTs.

### Gadolinium inhibition and the analogy to mechanosensitive channels

Further support for the nanopore hypothesis comes from our recent findings that Gadolinium (Gd^3+^)—a trivalent lanthanide known to block mechanosensitive and voltage-gated ion channels—also abolishes MT oscillations [11]. Gd^3+^ interacts strongly with negatively charged residues such as glutamate and aspartate, which are abundant at the nanopore lining of the αβ-tubulin interfaces. These residues create a localized electrostatic environment analogous to the selectivity filters of conventional ion channels. Gd^3+^ binding to these sites likely neutralizes the pore charge and prevents ionic exchange, thus collapsing the oscillatory regime. The inhibitory effect of Gd^3+^ was concentration-dependent and reversible, consistent with classical pore-blocking behavior observed in other biological ion channels.

This observation provides strong mechanistic evidence that the nanopores behave as functional ionic channels, whose gating can be modulated or blocked by specific ligands. The resemblance to the action of Gd^3+^ on mechanosensitive and stretch-activated channels (such as Piezo or TRP family members) reinforces the notion that nanopores act as biophysical transduction elements, responding to local electric or mechanical perturbations.

### Integration with the nonlinear electrokinetic model

The theoretical model presented here describes the MT as two coupled nonlinear transmission lines—representing the luminal and surface ionic pathways— interconnected by voltage-dependent nanopores. The coupling allows the emergence of leapfrogging solitons, where energy alternates between the inner and outer currents, giving rise to self-sustained oscillations. The experimental inhibition of oscillations by Taxol and Gd^3+^ provides direct validation of this model: both agents selectively block the interfacial nanopores, thereby disrupting the coupling that sustains the oscillatory regime. In this context, the oscillations are not merely emergent artifacts but represent an intrinsic electrodynamic property of the MT lattice that depends critically on nanopore-mediated ionic exchange.

### Functional implications in cellular signaling

From a physiological standpoint, these findings suggest that MTs may serve as active elements of intracellular signaling networks, rather than passive cytoskeletal supports. Oscillatory ionic currents could convey information in a frequency-encoded manner, coordinating processes such as vesicular transport, ciliary beating, or synaptic signaling. The sensitivity of MT oscillations to pharmacological agents like Taxol and ionic modifiers like Gd^3+^ also implies that cells may regulate cytoskeletal electrical activity via endogenous ionic or mechanical modulators. Given that MT oscillation frequencies overlap (among others, Scarinci et al., 2023[6]) with the gamma range of neuronal activity (30–100 Hz), it is plausible that MT-based electrodynamics contribute to synchronization phenomena underlying higher-order brain functions.

### Conclusion

In summary, the combined theoretical and experimental evidence underscores the central role of nanopores in the electrical activity of MTs. Pharmacological inhibition by Taxol and Gd^3+^ demonstrates that these pores are not passive structural defects but functional nanogates whose state determines the oscillatory and amplifying properties of the polymer. Their coupling to the surrounding ionic environment enables MTs to act as biological transistors, mediating nonlinear amplification and signal propagation at the nanoscale.

By situating these phenomena within the broader context of cell signaling, our findings extend the frontier of cytoskeletal biophysics, positioning MTs as active bioelectronic elements with potential relevance for cellular communication, neuronal dynamics, and emergent quantum–electrodynamic phenomena in biology. Rather than relying solely on diffusion-limited processes, cells could harness frequency-specific oscillatory signals along cytoskeletal filaments to regulate local biochemical events, synchronize activity across compartments, and potentially contribute to higher-order integrative processes in excitable tissues such as the brain. This biological framing underscores the broader significance of our model, making it accessible not only to physicists and engineers but also to experimental biologists interested in cytoskeletal dynamics and cell communication.

## Methods

We introduce a novel hollow cylindrical biomolecule model for tubulins (Fig. 10) and a two coupled electrical transmission lines prototype model (Fig. 11) to characterize the ionic electrical conductivity of MTs in intracellular aqueous electrolyte solutions. This approach expands upon the transmission line framework developed for our actin filament model [17].

**Figure 10.**
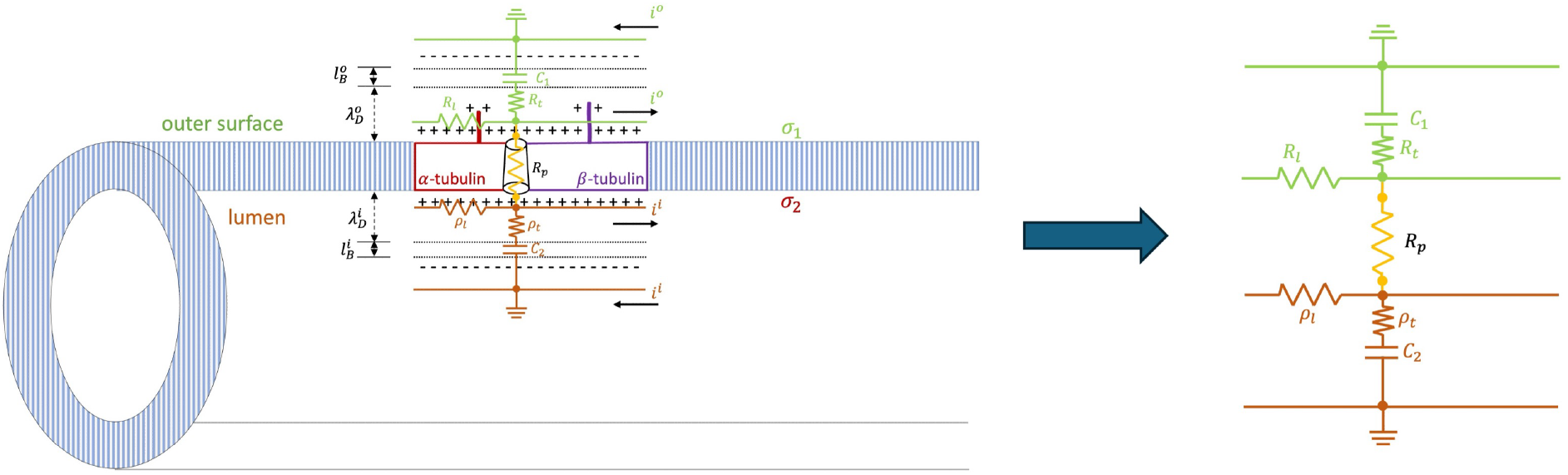
Multi-scale electrokinetic model for microtubules. It represents the outer and inner EDLs as coupled transmission lines (green and brown) connected through nanopores in the microtubule wall (yellow), modeling tubulin as a hollow cylindrical structure.

**Figure 11.**
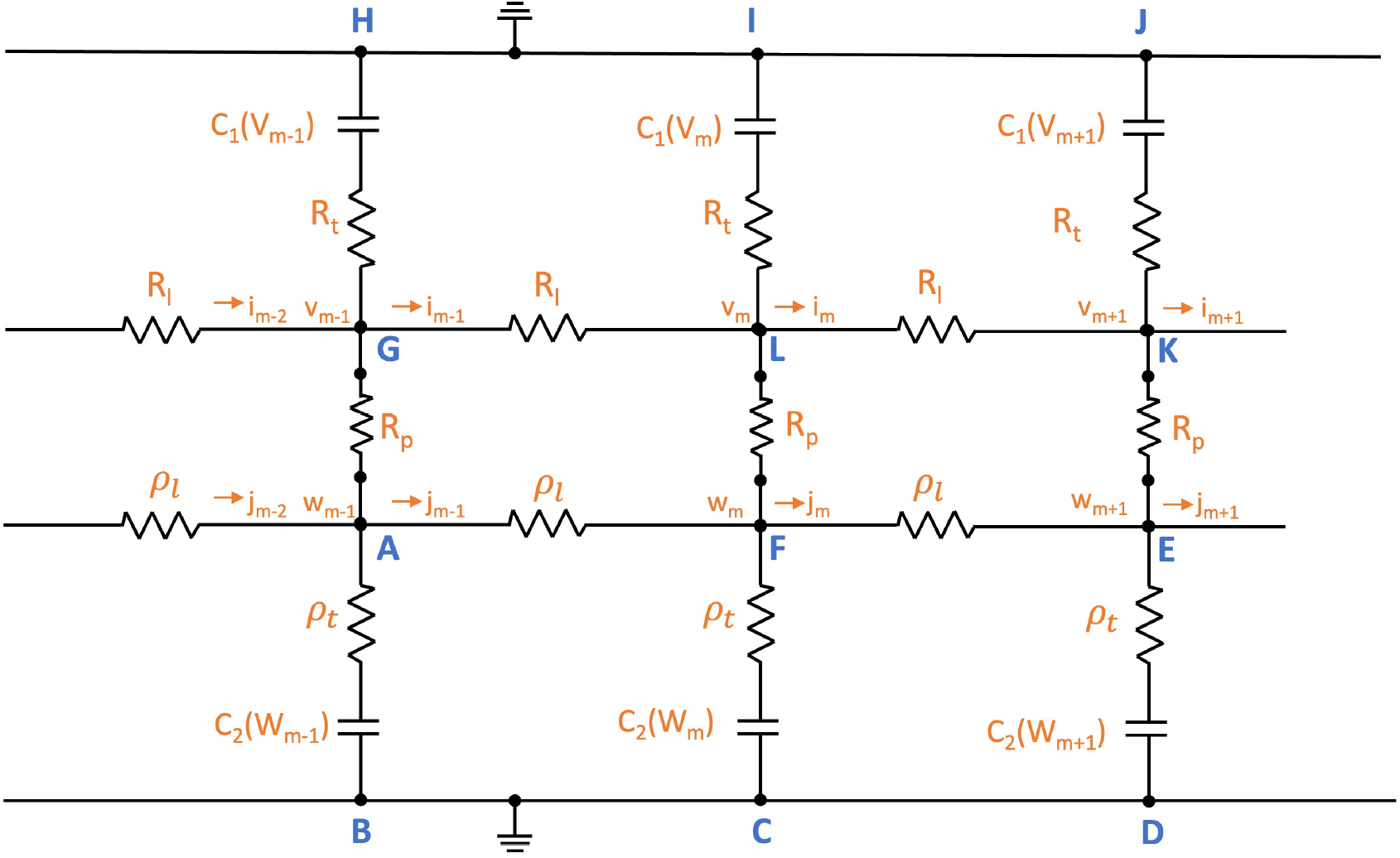
Ladder transmission lines model. The electric circuit unit cell represents an elementary ring of tubulin dimers shown in Fig. 10.

We retrieved the MT molecular structure information using the cryo-electron microscopy model 3j2u posted on the protein data bank [27]. It provided a detailed molecular characterization, including the amino acid sequence and the number and type of residues exposed to the electrolyte. This uncharged molecular structure in pdb format was uploaded into CSDFTS software [28] to assign atomic charges and sizes, add hydrogens, optimize the hydrogen bonding network, and renormalize atomic charges of the residues exposed to the surface due to pH effects (protonation/deprotonation process). Molecular structure analysis indicated that the single MT resembles a long, hollow cylindrical biomolecule with an outer diameter of approximately 25 nm, a wall thickness of about 5 nm, and a hollow central core measuring around 15 nm in diameter. The number of protofilaments in MTs typically ranges from 10 to 15 in-vitro, with 13 being the most common (Fig. 12).

**Figure 12.**
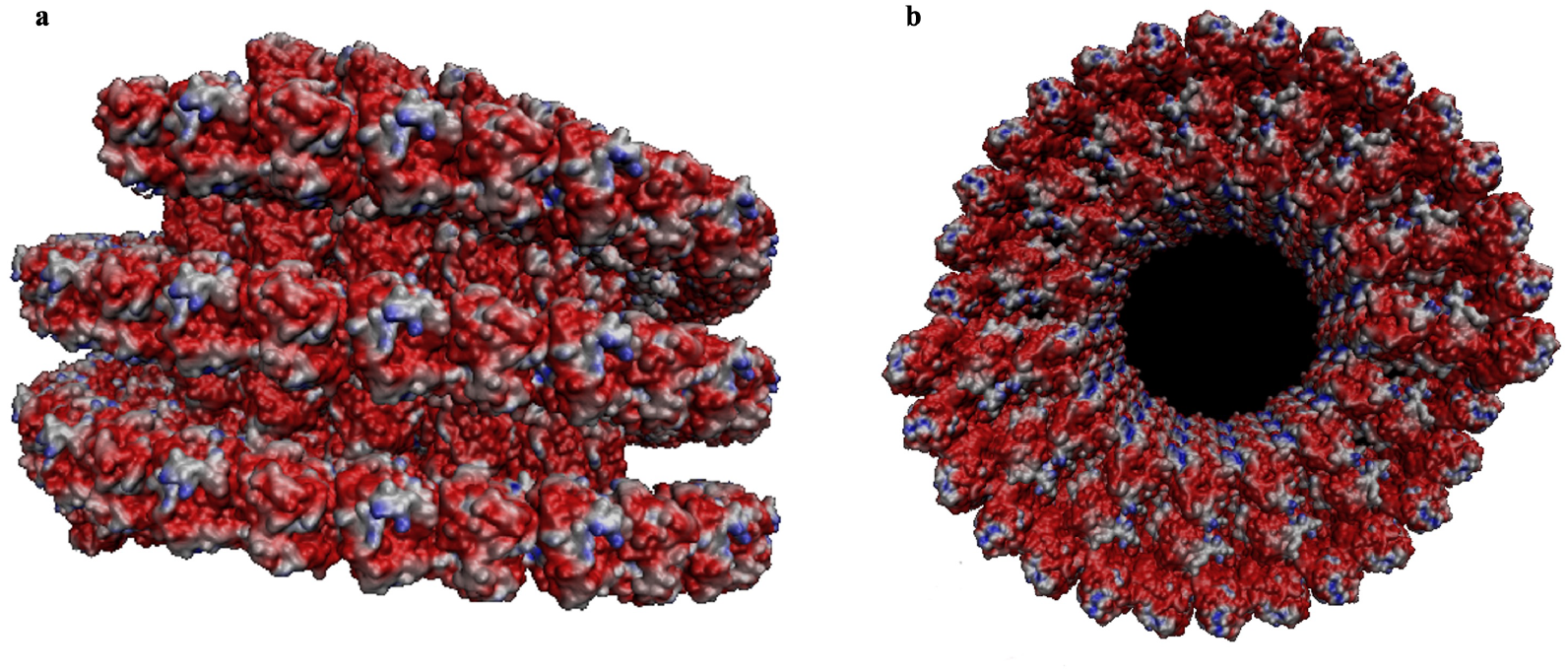
MT molecular structure and surface charge distribution. (a) Longitudinal view. (b) Cross-section view. Cryo-electron microscopy data (PDB ID: 3j2u) were used to model the microtubule structure and surface charge distribution. Red indicates negative charges, blue positive charges, and white neutral regions.

In this context, we modeled MTs as polyelectrolytes with negative charges uniformly distributed on the tubulin surface with an approximate charge ratio of 2:1 between the outer and inner surfaces [9]. We characterized each transmission line’s electrical and conductive properties using the hollow cylindrical biomolecule model for tubulin dimers, Ohm’s law, Navier-Stokes, and Poisson’s theories. Specific considerations on the outer and inner transmission lines and their coupling interaction via nanopores are provided below. More details on these calculations can be found in the supplementary document.

### Outer Transmission Line Model

Here, we highlight the modifications and extensions done on a single transmission line model for actin filaments [17] to characterize the outer transmission lines for MTs. We replace the elementary circuit unit of an actin monomer with an elementary ring of 13 dimers of a radius of 12.5 *nm* and length of *l* = 8 *nm*. Similarly, the EDL capacitor accounts for a nonlinear condensed ionic cloud in the radial direction on the MT that changes with applied voltage and contributes to the nonlinearity of the electrical impulse. Unlike G-actin monomers, tubulins have negatively charged C-terminal tubulin tails (TTs) that extend 4–5 *nm* from the microtubule surface. Since TTs undergo shrinking [8, 9, 16] and thermal oscillations [21], they contribute to MT nonlinearity. These properties are considered using the nonlinear charge condensation

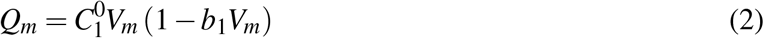

where *V*_*m*_ represents the outer unit circuit voltage, and 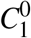 stands for the linear part of the capacitance. The nonlinear capacitance calculations performed for G-actin proteins [17] cannot be extended to tubulins because they are valid for solid cylinders only. Instead, here we solved the linearized Poisson Boltzmann equation for the electric potential of an infinite long tubular cylinder to get the value of the linear capacitance for the outer line 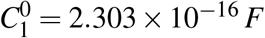 (see supplementary document for more details). In this model, the small nonlinear parameter *b*_1_ is unknown since it represents the change in capacitance with voltage, coming from the outer condensed ionic cloud and TTs. It is assumed to be a free parameter of the model, and its value is set equal to *b*_1_ = 0.06 *V* ^−1^, which is small compared to the voltage stimulus range considered in these studies.

Another MT feature considered in this model is the dispersion of the electrical impulse generated by the structural periodicity manifest in the tubulin dimers arrangement. Meanwhile, the longitudinal and transversal resistors account for the losses in the transmission media. By combining the electric potential solution for a tubular cylinder and the electrokinetic ionic current expressions coming from Ohm’s law and Poisson’s and Navier-Stokes’ equations, we calculated the longitudinal resistance for the outer line, yielding a value of *R*_*l*_ = 5.845 × 10^6^ Ω (see supplementary document for more details). In contrast, the transversal resistance calculations performed for G-actin proteins [17] cannot be extended to tubulins because they do not account for MT nanopores and TTs, which is not an obstacle for single transmission line models [9] since the transversal resistance is supposed to be very high. The leakage of ionic currents through the depleted layer can be ignored. However, in the leapfrogging model of coupled two transmission lines, current leakages through the depleted layer and nanopore significantly influence soliton oscillations. Thus, we considered a high value for the outer transversal resistance *R*_*t*_ = 7.792 × 10^14^ Ω, which is in the same order as the pore resistance. Additionally, we set the outer impedance to *Z*_1_ = 1.862 × 10^11^ Ω, and the inductance is so small (of the order of ∼10^−16^ *H*) compared to the other impedance components that we ignored its effect [8, 9, 16].

### Inner Transmission Line Model

Figure 11 shows that the inner electrical transmission line mirrors the outer transmission line described above but has different (asymmetric) electric circuit parameter values. Here, we remark on the main changes to the outer transmission line calculations to characterize the electric circuit for the lumen. We consider a negatively charged inner surface with a surface charge density ratio of 1:4 to the outer surface [29]. We define the nonlinear capacitance of the inner line as

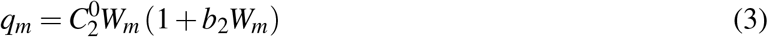

where *W*_*m*_ represents the inner unit circuit voltage, and 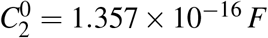 represents the inner linear capacitance. We noticed that 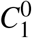 is greater than 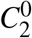 since the outer condensed ionic cloud thickness is four times greater than the inner one. Additionally, *b*_2_ denotes a small lumen nonlinearity parameter, which we set equal to *b*_2_ = 0.03 *V* ^−1^, a lower value than the outer one because there are no TTs in the former. Another essential difference between the outer and inner lines is the nonlinear term sign, which accounts for the opposite polarities of the voltage-dependent capacitors on the two lines [23]. The calculations for the longitudinal resistance of the inner line yielded *ρ*_*l*_ = 1.810 × 10^7^ Ω. Additionally, we estimated the inner transversal resistance and impedance by *ρ*_*t*_ = 3.779 × 10^14^ Ω and *Z*_2_ = 3.126 × 10^11^ Ω, respectively.

### Incorporating Time-Variant Pore Conductance into the Transistor Model of Microtubule Oscillations

MTs exhibit electrical properties that suggest their role as dynamic conductors of ionic currents. Their structure, characterized by highly charged surfaces and periodic nanopores, enables them to function as biological transmission lines. However, beyond this passive conductive role, experimental observations indicate that MTs amplify electrical signals [15, 24], sustain oscillatory dynamics, and exhibit voltage-dependent behavior [2, 3, 5]. These features suggest a functional analogy to transistors, where the nanopores act as gates regulating the flow of ions in response to external stimuli. In our model, the voltage-dependent permeability of α–β interdimer nanopores serves as the biophysical basis for this gate-like modulation.

In classical transistors, a small control signal at the gate modulates a larger current between the source and drain terminals. A similar mechanism is present in MTs, where nanopores’ periodic opening and closing control ionic flow across the structure, leading to oscillatory charge dynamics that can be modeled in various transistor frameworks. In a FET model, the nanopores act as gate-controlled conduits for ionic currents, where the electrostatic potential near the pore entrance modulates the conductance. This dynamic regulation of the pore conductance resembles the behavior of a voltage-gated FET, where the local electric field influences the density and mobility of charge carriers. In this case, the drain-source current can be expressed as

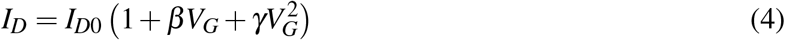

where *I*_*D*0_ is the baseline drain current, *V*_*G*_ is the gate voltage (nanopore potential), β is the gate modulation factor (linear term), and γ accounts for higher-order nonlinearities in conductance. The dynamic modulation of ionic charge carrier mobility could also be described as

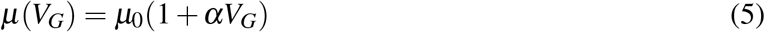

where *µ*_0_ is the baseline charge carrier mobility and α describes the dependence of mobility on gate voltage.

In JFETs, a depletion region controls the current. Similarly, in MTs, counterion condensation and depletion within the nanopores modulate the ionic flow, creating a time-dependent gate potential that governs oscillatory conduction. The conductance can be modeled as

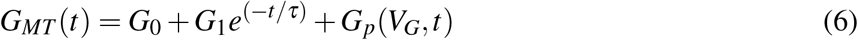

where *G*_0_ is the baseline conductance, *G*_1_ represents transient gating effects that decay over time with characteristic time constant τ, and *G*_*p*_(*V*_*G*_, *t*) is the voltage- and time-dependent pore conductance.

In a Bipolar Junction Transistor (BJT) analogy, nanopores serve as the base region, where small variations in ionic concentration similar to the base current lead to amplified changes in longitudinal ionic currents (collector-emitter current). The current amplification can be described as

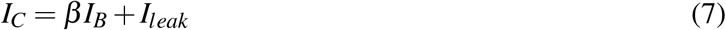

where *I*_*C*_ is the ionic collector current, β is the current gain, *I*_*B*_ is the base current from nanopore dynamics, and *I*_*leak*_ accounts for ionic leakage through the MT wall.

The amplifying function of the MT system can be further characterized by defining an effective β parameter for ion transport

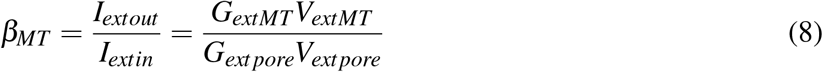

where *G*_*MT*_ and *V*_*MT*_ represent the total microtubule conductance and voltage, while *G*_*pore*_ and *V*_*pore*_ capture the localized gating effects of the nanopores.

This amplification mechanism suggests that small local perturbations in ionic concentrations at the nanopores could lead to significant long-range signal propagation, similar to transistor-based amplification in electronic circuits.

### Coupling to Leapfrogging Transmission Lines

We consider that the outer and inner electrical transmission lines are coupled through nanopores. The nanopores are permeable to counterions from the ionic clouds on the inner and outer surfaces [7, 9] and impermeable to anions [7]. Thus, counterions are exchanged between the inner and outer transmission lines through the nanopores. Voltage-clamp experimental data on MTs show that the oscillating current frequency (∼39 Hz) does not depend on the voltage stimulus [2, 3, 5]. This result implies that nanopores can be characterized by a voltage-dependent nonlinear resistance *R*_*p*_(*V*) [16], directly influencing the coupled soliton dynamics. This introduces a self-regulatory mechanism where the interplay between ion accumulation and depletion cycles at the pores creates a time-dependent gating effect. The result is a leapfrogging behavior, where ionic solitons continuously overtake each other, similar to the oscillatory charge redistribution in a transistor circuit [22, 23].

To determine *R*_*p*_(*V*), we kept all the circuit parameters in Fig. 4 fixed, and adjusted pore resistance values in the input voltage range between 0.01 V and 0.1 V to reproduce the experimental oscillation frequency. Figure 13-a displays our results. The corresponding I-V curve was computed using a reversal potential of 0.005 V to reflect the slight asymmetry between the intra- and extra-microtubule ion concentrations. The data, shown in Fig. 13-b, evidence a good fitting with 2S3B energy model for ion permeation through the nanopore [2, 30] given by the following expression

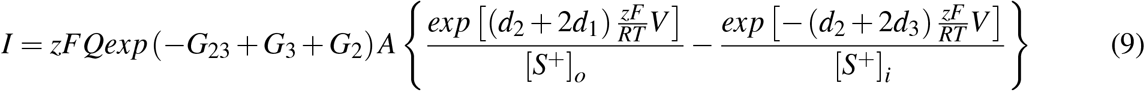

where [*S*^+^]_*o*_ and [*S*^+^]_*i*_ are the concentrations of permeable ion in the MT outer and intra environment, respectively, *V* is the potential, *F* is the Faraday constant, *R* is the universal gas constant, *T* stands for the temperature, *z* represents the ion valence, and *Q* is a prefactor encompassing the transition rate constants between pore states. Also, *G*_23_ is the peak energy, *G*_2_ and *G*_3_ are the well energies, and *d*_1_, *d*_2_, and *d*_3_ are the three electrical distances. The optimal parameter values obtained in our fitting approach are *A* = 2.16, *G*_2_*/G*_3_ = −0.67, *G*_23_ = −0.70, *d*_1_ = 0.03, *d*_2_ = 0.04, and *d*_3_ = 0.43. These values are qualitatively similar to the values obtained in fitting voltage clamp experimental data [2].

**Figure 13.**
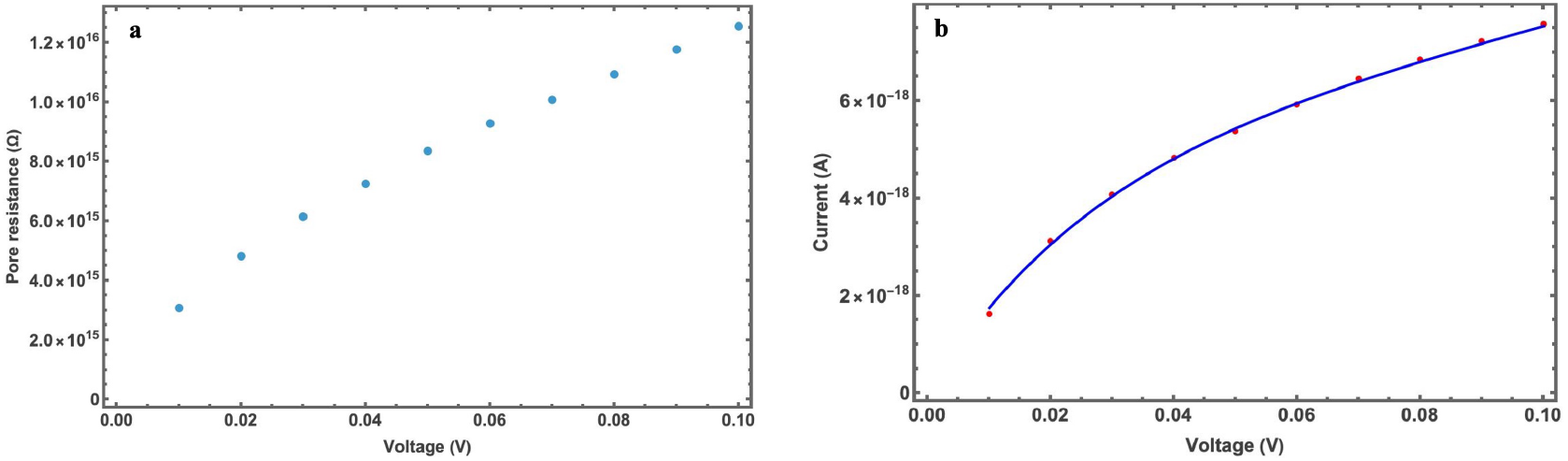
Voltage-dependent nanopore resistance model. a) The pore resistance *R*_*p*_(*V*) is calculated for input voltages ranging from 0.01 V to 0.1 V, keeping the other circuit parameters fixed. b) The corresponding current-volrage (I-V) curve shows outward rectification. Red dots represent the calculated data points, and the blue curve corresponds to the fit using the 2S3B energy model.

### Transmission Lines Model Solution

We solved the coupled electrical transmission lines following the procedure introduced in our model for actin filaments [17]. Applying Kirchoff’s rules to the unit cell number *m*, we obtained equations S.25 and S.26 provided in the supplementary document (see Fig. 11). We used the notation 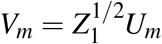 and 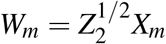 for the outer and inner voltage lines. Since the circuit unit length is much less than the MT length (i.e., the length of the transmission line), voltages and currents in the transmission lines change gradually from one circuit unit to the next. This allowed us to use the continuum approximations *U*_*m*_ (*t*) ≃ *U* (*x, t*) and *X*_*m*_ (*t*) ≃ *X* (*x, t*)J, and to apply the transformations to the space and time coordinates 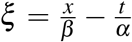 and 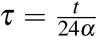, with 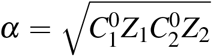 and *β* = 2*l* (*l* is the length of the dimer, e.g. 8 nm). Finally, we rescaled the transmission line voltages 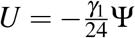 and 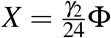 in equations S.31 and S.32 in the supplementary document, with 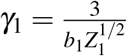 and 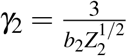, to obtain the following two master coupled perturbative KdV equations

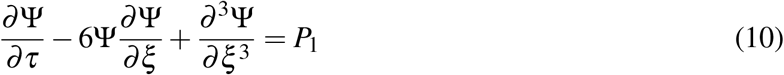

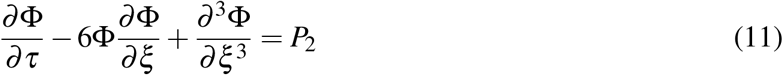

where,

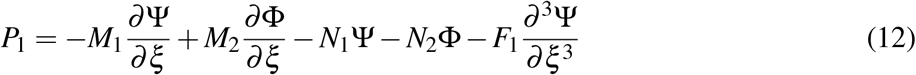

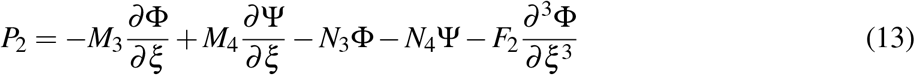

Equations 10 and 11 describe the nonlinear dynamics of voltage perturbations along the outer (Ψ) and inner (Φ) ionic conduction paths of the MT. These variables represent normalized voltage amplitudes propagating along the outer surface and luminal pathways, respectively. The coupling through nanopores introduces nonlinear terms that enables sustained oscillatory solutions.

Labels 1 and 2 on P represent the outer and inner transmission lines. The expressions for the parameters *N*’s, *M’s*, and *F’s* in equations 12 and 13 are linked to the electric circuit parameters through the equations S.33 to S.44 in the supplementary document. These parameters are computed from the microtubule’s electrical and geometrical properties. It is worth noting that *P*_1_ and *P*_2_ combine coupling and self-perturbation terms. The first terms on the left of equations 10 and 11 resemble the time-dependent term in Fick’s diffusion law. The second and third terms represent the nonlinearity and dispersive contributions arising from the condensed ion cloud in the EDL and the diffuse spreading of ions along the MT, respectively. On the other hand, the first and second terms to the right in equations 12 and 13 represent the electric fields generated by the ions in the outer(inner) and inner(outer) EDLs and resemble the potential gradient term in the Nernst-Plank electrodiffusion equation. The third and fourth terms to the right represent the damping perturbations.

To solve the perturbative KdV equations 10 and 11, we followed a similar procedure introduced in our actin filament article [17] and the leapfrogging articles [22, 23]. The exact soliton solutions for *P*_1_ = *P*_2_ = 0 correspond to the homogenous KdV equations given by

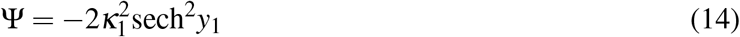

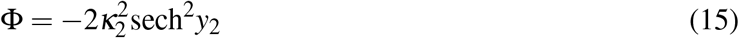

where *y*_1_ = *κ*_1_ (*ξ* − *θ*_1_) and 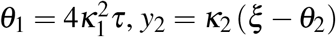 and 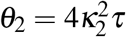.

According to the adiabatic perturbation theory [26], the perturbations *P*_1_ and *P*_2_ modify these homogenous solutions, which is accounted for by introducing time-dependence on the amplitudes *κ*_*i*_ → *κ*_*i*_ (*τ*) and solving the following integrodifferential equations

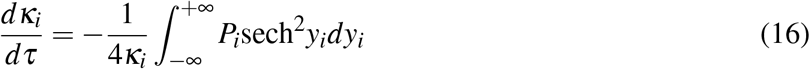

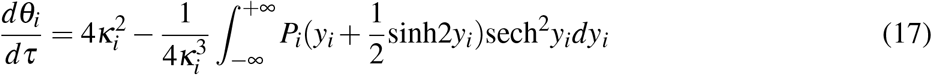

Substitution of equations 12, 13, 14, and 15 into equations 16 and 17 yielded the four coupled firstorder asymmetric ordinary differential equations S.47, S.50, S.57, and S.58. We solved these equations numerically using Mathematica software (see the supplementary document).

### Analytical Solution for Phase and Amplitude Dynamics of Coupled Solitons

We numerically solved the coupled ODE system–equations for multiple values of the involved parameters and concluded that the parameters *M*_1_, *M*_3_, *N*_2_, *N*_4_, *F*_1_, and *F*_2_ change the soliton amplitudes but they weakly affect their oscillation frequency. Thus, we neglected these parameters and linearized the coupled ODE system–equations for small Δ*θ* and Δ*κ* to obtain equations S.59–S.64. These coupled equations were solved in the supplementary document to obtain an approximate analytical expression for the soliton oscillation frequency. In doing so, we got the following damped oscillatory equation

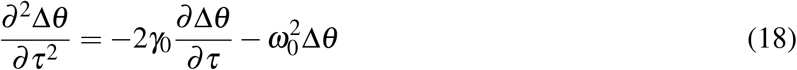

Where 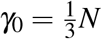, and 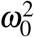 is given by Eq. 1. We considered *κ*_1_ = *κ* + *λ*_1_, *κ*_2_ = *κ* + *λ*_2_, Δ*λ* = *λ*_1_ − *λ*_2_, Δ*θ* = *θ*_1_ − *θ*_2_, and *N*_1_ = *N*_3_ = *N*. Equation 1 provides the analytical expression for the dominant oscillation frequency *ω*_0_.

Using the initial conditions Δ*θ* (*τ* = 0) = Δ*θ*_0_ and 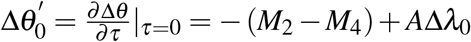, we get the following analytic solution for the soliton phase difference

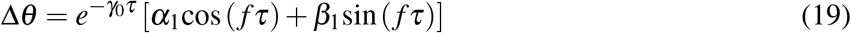

Where 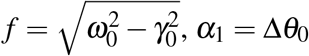, and 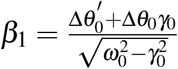.

Similarly, the following damped oscillatory equation for the amplitude difference was obtained

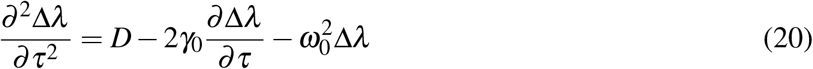

Where 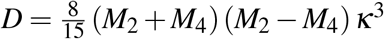. Using 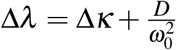, we can rewrite the previous equation as follows

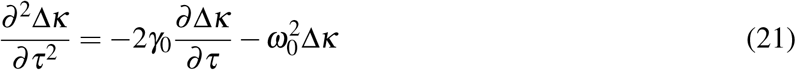

As a result, the analytic solution can be expressed as

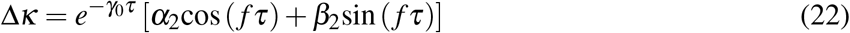

where *α*_2_ = Δ*κ*_0_ and 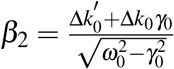. The initial conditions are given by 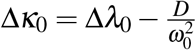 and 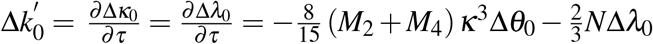.

Equations 18–22 describe the evolution of amplitude (Δ*κ*) and phase (Δ*θ*) differences between inner and outer solitons under dissipative conditions. The decay constant *γ*_0_ reflects dissipative losses in the ionic double layer, while *ω*_0_ is controlled by nanopore coupling.

## Acknowledgments

M. Mohsin and Dr. M. Marucho thank Dr. Dikande Alain Moise and Dr. Nkongho Achere Akem for sharing calculations of the leapfrogging approach and Dr. Vadim Warshavsky for his contribution to the Poisson’s equation solution for MTs.

## Author Contributions

M. Marucho designed the research; M. Mohsin and M. Marucho developed the mathematical model and performed simulations; M. Mohsin prepared all the figures corresponding to the model and numerical results; H. Cantiello and M. d.R. Cantero provided experimental validation and prepared Fig. 1; M. Mohsin and M. Marucho wrote the main manuscript text; all authors reviewed and edited the manuscript.

## Data Availability

The datasets and Mathematica notebooks used and/or analysed during the current study are available from the corresponding author’s GitHub repository website https://github.com/MarceloMarucho/EOMT/.

## Additional Information

Additional information is provided in the supplementary document.

