## Supplemental document for "Electrical Oscillations in Microtubules"

### Poisson's equation solution, and capacitance and resistance calculations

In our model, the microtubule occupies the volume between two infinitely long coaxial cylinders with radii $R_1$ and $R_2$, $R_1 < R_2$ (Figure 8 in the main text). The surface charge densities of the inner and the outer cylinder surfaces are $\sigma_1$ and $\sigma_2$, correspondingly. The dielectric permittivity of the area between the cylinders $R_1 \leq r \leq R_2$ (i.e., inside the microtubule) is $\varepsilon_2$. The electrolyte solution fills the space outside the microtubule with $r \leq R_1$ and $r \geq R_2$. The dielectric permittivity of the electrolyte solution is $\varepsilon$.To find the electrostatic potential $\varphi(r)$ in the whole system, we write the set of Poisson's equations for the system

$$\begin{cases} \Delta\varphi_1(r) = -\frac{\rho_1(r)}{\varepsilon}, & r \leq R_1 \\ \Delta\varphi_2(r) = 0, & R_1 \leq r \leq R_2 \\ \Delta\varphi_3(r) = -\frac{\rho_3(r)}{\varepsilon}, & R_2 \leq r \end{cases} \tag{S.1}$$

where $\rho(r)$ stands for the electric charge density in the electrolyte solution.

The solutions of the corresponding linearized Poisson Boltzmann equations are

$$\begin{cases} \varphi_1(r) = AI_0\left(\frac{r}{\lambda_D}\right), & r \leq R_1 \\ \varphi_2(r) = B\ln\frac{r}{R_1} + C, & R_1 \leq r \leq R_2 \\ \varphi_3(r) = DK_0\left(\frac{r}{\lambda_D}\right), & R_2 \leq r \end{cases} \tag{S.2}$$

Using the boundary conditions, we obtained the following expressions for the coefficients $A$, $B$, $C$, and $D$

$$\begin{gathered} A = C\frac{1}{a_1} \\ B = \frac{a_2}{a_1}C - a_3 \\ C = \frac{\left(\frac{a_5a_8}{a_6} + a_3a_4 + \frac{a_3a_5a_7}{a_6}\right)}{\left(\frac{a_2a_4}{a_1} + \frac{a_5a_7a_2}{a_6a_1} + 1\right)} \\ D = -\frac{a_2a_7}{a_1a_6}C + \frac{a_8}{a_6} + \frac{a_3a_7}{a_6} \end{gathered} \tag{S.3}$$

with,

$$a_1 = I_0\left(\frac{R_1}{\lambda_D}\right) \tag{S.4}$$

$$a_2 = \frac{\varepsilon}{\varepsilon_2}\frac{R_1}{\lambda_D}I_1\left(\frac{R_1}{\lambda_D}\right) \tag{S.5}$$

$$a_3 = \frac{\sigma_1 R_1}{\varepsilon_2} \tag{S.6}$$

$$a_4 = \ln\frac{R_2}{R_1} \tag{S.7}$$

$$a_5 = K_0\left(\frac{R_2}{\lambda_D}\right) \tag{S.8}$$

$$a_6 = K_1\left(\frac{R_2}{\lambda_D}\right) \tag{S.9}$$

$$a_7 = \frac{\varepsilon_2}{\varepsilon}\frac{\lambda_D}{R_2} \tag{S.10}$$

$$a_8 = \frac{\sigma_2 \lambda_D}{\varepsilon} \tag{S.11}$$

where $\lambda_D = \sqrt{\frac{\varepsilon RT}{F^2 \sum_i z_i^2 c_{i,0}}}$ is the Debye length.

Using the Boltzmann distribution and the previous solution for the electric pontetial, the ion density distributions can be calculated as follows

$$\begin{cases} c_{i,1} = c_{i,0}\left[1 - \frac{z_i F}{RT}\varphi_1(r)\right], & r \leq R_1 \\ \\ c_{i,3} = c_{i,0}\left[1 - \frac{z_i F}{RT}\varphi_3(r)\right], & R_2 \leq r \end{cases} \tag{S.12}$$

where $c_{i,0}$ and $z_i$ are the bulk concentration and valence of species $i$, respectively, R is the gas constant, T the electrolyte temperature, and F the Faraday's constant.

The corresponding expression for the electrolyte conductivities are

$$\begin{cases} k_1 = F^2 \sum_i z_i^2 u_i c_{i,1}, & r \leq R_1 \\ \\ k_3 = F^2 \sum_i z_i^2 u_i c_{i,3}, & R_2 \leq r \end{cases} \tag{S.13}$$

and the total charge density distributions are

$$
\begin{cases}
\rho_1 = F\sum_i z_i c_{i,1}, & r \le R_1 \\
\\
\rho_3 = F\sum_i z_i c_{i,3}, & R_2 \le r
\end{cases}
\tag{S.14}
$$

Combining Poisson and the Navier-Stokes' equations, we obtained the following expressions for the axial velocity profiles

$$
\begin{cases}
v_{z,1} = \frac{\varepsilon E_z}{\mu}\left[\varphi_1(r) - \varphi_1(R_1)\right], & r \le R_1 \\
\\
v_{z,3} = \frac{\varepsilon E_z}{\mu}\left[\varphi_3(r) - \varphi_3(R_2)\right], & R_2 \le r
\end{cases}
\tag{S.15}
$$

where $\mu$ represents the viscosity parameter, $E_z = V/\ell$, $V$ is the input voltage, and $\ell$ the tubulin size.

Using the expressions S.2-S.15, we obtained the following expression for the longitudinal currents

$$
\begin{cases}
\frac{I_{l,1}}{2\pi} = E_z\int_{R_1-l_B}^{R_1} rk_1(r)\,dr + \int_{R_1-l_B}^{R_1} rv_{z,1}\rho_1(r)\,dr, & r \le R_1 \\
\\
\frac{I_{l,3}}{2\pi} = E_z\int_{R_2}^{R_2+l_B} rk_3(r)\,dr + \int_{R_2}^{R_2+l_B} rv_{z,3}\rho_3(r)\,dr, & R_2 \le r
\end{cases}
\tag{S.16}
$$

where $l_B = \frac{e^2}{4\pi\varepsilon k_B T}$ is the Bjerrum length, and $k_B$ the Boltzmann constant.

We used Ohm's law, which relates the axial voltage drop $\Delta V$ and the total electric current $I_l$ as $R_l = |\frac{\Delta V}{I_l}|$, to obtain the expression for longitudinal resistances for the outer and inner surfaces of the microtubule.

Finally, we plot $\frac{d\varphi}{d\sigma}$ to calculate the linear capacitance parameters $C_1$ and $C_2$ from the slope for both inner and outer MT surfaces, respectively.

### Coupled KdV equations

$\frac{\partial Q_m}{\partial t}$ and $\frac{\partial q_m}{\partial t}$ represent the currents in the cell $m$ across the capacitors $C_1$ and $C_2$, respectively, while $Q_m$ and $q_m$ are given by the equations 1 and 2 in the main text. Applying the current conservation law in the nodes L and F between the cells $m-1$ and $m$, provides the following equations for the currents (Figure 9 in the main text)

$$
i_{m-1} - i_m = \left(1 + \frac{R_t}{R_p}\right)C_1^0\left(\frac{\partial V_m}{\partial t} - 2b_1 V_m \frac{\partial V_m}{\partial t}\right) + \frac{V_m}{R_p} - \frac{\rho_t}{R_p}C_2^0\left(\frac{\partial W_m}{\partial t} + 2b_2 W_m \frac{\partial W_m}{\partial t}\right) - \frac{W_m}{R_p} \tag{S.17}
$$

$$
j_{m-1} - j_m = \left(1 + \frac{\rho_t}{R_p}\right)C_2^0\left(\frac{\partial W_m}{\partial t} + 2b_2 W_m \frac{\partial W_m}{\partial t}\right) + \frac{W_m}{R_p} - \frac{R_t}{R_p}C_1^0\left(\frac{\partial V_m}{\partial t} - 2b_1 V_m \frac{\partial V_m}{\partial t}\right) - \frac{V_m}{R_p} \tag{S.18}
$$

Additionally, using the Kirchoff's law to along the loop, we obtain

$$\left(1+\frac{R_t}{R_p}\right) i_m.R_l = R_t(i_{m-1}-2i_m+i_{m+1})+(V_m-V_{m+1})+\frac{R_t}{R_p}j_m\rho_l \tag{S.19}$$

$$\left(1+\frac{\rho_t}{R_p}\right) j_m.\rho_l = \rho_t(j_{m-1}-2j_m+j_{m+1})+(W_m-W_{m+1})+\frac{\rho_t}{R_p}i_mR_l \tag{S.20}$$

Using

$$i_m = Z_1^{-1/2}U_m; \qquad Z_1^{1/2}U_m = V_m$$

$$j_m = Z_2^{-1/2}X_m; \qquad Z_2^{1/2}X_m = W_m$$

equation S.17 becomes

$$U_{m-1}-U_m = \left(1+\frac{R_t}{R_p}\right)C_1^0Z_1\left(\frac{\partial U_m}{\partial t}-2b_1Z_1^{1/2}U_m\frac{\partial U_m}{\partial t}\right)+\frac{Z_1U_m}{R_p}$$
$$-\frac{\rho_t}{R_p}C_2^0Z_1^{1/2}Z_2^{1/2}\left(\frac{\partial X_m}{\partial t}+2b_2Z_2^{1/2}X_m\frac{\partial X_m}{\partial t}\right)-\frac{Z_1^{1/2}Z_2^{1/2}X_m}{R_p} \tag{S.21}$$

equation S.18 becomes

$$X_{m-1}-X_m = \left(1+\frac{\rho_t}{R_p}\right)C_2^0Z_2\left(\frac{\partial X_m}{\partial t}+2b_2Z_2^{1/2}X_m\frac{\partial X_m}{\partial t}\right)+\frac{Z_2X_m}{R_p}$$
$$-\frac{R_t}{R_p}C_1^0Z_1^{1/2}Z_2^{1/2}\left(\frac{\partial U_m}{\partial t}-2b_1Z_1^{1/2}U_m\frac{\partial U_m}{\partial t}\right)-\frac{Z_1^{1/2}Z_2^{1/2}U_m}{R_p} \tag{S.22}$$

equation S.19 becomes

$$U_m-U_{m+1} = \frac{R_t}{R_t^{\text{equiv.}}}Z_1^{-1}U_mR_l - R_tZ_1^{-1}(U_{m-1}-2U_m+U_{m+1})-\frac{R_t}{R_p}Z_1^{-1/2}Z_2^{-1/2}X_m\rho_l \tag{S.23}$$

where,

$$\frac{1}{R_t^{\text{equiv.}}}=\frac{1}{R_p}+\frac{1}{R_t}$$

and, equation S.20 becomes

$$X_m - X_{m+1} = \frac{\rho_t}{\rho_t^{\text{equiv.}}} Z_2^{-1} X_m \rho_l - \rho_t Z_2^{-1}(X_{m-1} - 2X_m + X_{m+1}) - \frac{\rho_t}{R_p} Z_1^{-1/2} Z_2^{-1/2} U_m R_l \tag{S.24}$$

where

$$\frac{1}{\rho_t^{\text{equiv.}}} = \frac{1}{R_p} + \frac{1}{\rho_t}$$

Combining equations S.21 and S.23, we get

$$\begin{aligned} U_{m-1} - U_{m+1} = & \frac{R_t}{R_t^{\text{equiv.}}} C_1^0 Z_1 \left( \frac{\partial U_m}{\partial t} - 2b_1 Z_1^{1/2} U_m \frac{\partial U_m}{\partial t} \right) + \left( \frac{Z_1}{R_p} + \frac{R_t}{R_t^{\text{equiv.}}} R_l Z_1^{-1} \right) U_m \\ & - R_t Z_1^{-1}(U_{m-1} - 2U_m + U_{m+1}) - \frac{\rho_t}{R_p} C_2^0 Z_1^{1/2} Z_2^{1/2} \left( \frac{\partial X_m}{\partial t} + 2b_2 Z_2^{1/2} X_m \frac{\partial X_m}{\partial t} \right) \\ & - \frac{Z_1^{1/2} Z_2^{1/2}}{R_p} \left( 1 + \frac{R_t \rho_l}{Z_1 Z_2} \right) X_m \end{aligned} \tag{S.25}$$

and combining equations S.22 and S.24, we get

$$\begin{aligned} X_{m-1} - X_{m+1} = & \frac{\rho_t}{\rho_t^{\text{equiv.}}} C_2^0 Z_2 \left( \frac{\partial X_m}{\partial t} + 2b_2 Z_2^{1/2} X_m \frac{\partial X_m}{\partial t} \right) + \left( \frac{Z_2}{R_p} + \frac{\rho_t}{\rho_t^{\text{equiv.}}} \rho_l Z_2^{-1} \right) X_m \\ & - \rho_t Z_2^{-1}(X_{m-1} - 2X_m + X_{m+1}) - \frac{R_t}{R_p} C_1^0 Z_1^{1/2} Z_2^{1/2} \left( \frac{\partial U_m}{\partial t} - 2b_1 Z_1^{1/2} U_m \frac{\partial U_m}{\partial t} \right) \\ & - \frac{Z_1^{1/2} Z_2^{1/2}}{R_p} \left( 1 + \frac{\rho_t R_l}{Z_1 Z_2} \right) U_m \end{aligned} \tag{S.26}$$

In the continuum limit, we can write

$$U_m(t) \simeq U(x,t)$$

$$U_{m\pm1} = U(x \pm l, t) \simeq U \pm l \frac{\partial U}{\partial x} + \frac{l^2}{2} \frac{\partial^2 U}{\partial x^2} \pm \frac{l^3}{3!} \frac{\partial^3 U}{\partial x^3} + \cdots$$

$$U_{m-1} - 2U_m + U_{m+1} = l^2 \frac{\partial^2 U}{\partial x^2}$$

$$U_{m-1} - U_{m+1} = -2l \frac{\partial U}{\partial x} - \frac{l^3}{3} \frac{\partial^3 U}{\partial x^3}$$

and

$$X_m(t) \simeq X(x,t)$$

$$X_{m\pm1} = X(x \pm l, t) \simeq X \pm l\frac{\partial X}{\partial x} + \frac{l^2}{2}\frac{\partial^2 X}{\partial x^2} \pm \frac{l^3}{3!}\frac{\partial^3 X}{\partial x^3} + \cdots$$

$$X_{m-1} - 2X_m + X_{m+1} = l^2\frac{\partial^2 X}{\partial x^2}$$

$$X_{m-1} - X_{m+1} = -2l\frac{\partial X}{\partial x} - \frac{l^3}{3}\frac{\partial^3 X}{\partial x^3}$$

Therefore, equation S.25 becomes

$$\frac{R_t}{R_t^{\text{equiv.}}}C_1^0 Z_1\frac{\partial U}{\partial t} - \frac{R_t}{R_t^{\text{equiv.}}}2b_1 Z_1^{3/2}C_1^0 U\frac{\partial U}{\partial t} + \left(\frac{Z_1}{R_p} + \frac{R_t}{R_t^{\text{equiv.}}}R_l Z_1^{-1}\right)U - R_t Z_1^{-1}l^2\frac{\partial^2 U}{\partial x^2} + 2l\frac{\partial U}{\partial x}$$
$$+\frac{l^3}{3}\frac{\partial^3 U}{\partial x^3} = \frac{\rho_t}{R_p}C_2^0 Z_1^{1/2}Z_2^{1/2}\left(\frac{\partial X}{\partial t} + 2b_2 Z_2^{1/2}X\frac{\partial X}{\partial t}\right) + \frac{Z_1^{1/2}Z_2^{1/2}}{R_p}\left(1 + \frac{R_t\rho_l}{Z_1 Z_2}\right)X \quad \text{(S.27)}$$

and equation S.26 becomes,

$$\frac{\rho_t}{\rho_t^{\text{equiv.}}}C_2^0 Z_2\frac{\partial X}{\partial t} + \frac{\rho_t}{\rho_t^{\text{equiv.}}}2b_2 Z_2^{3/2}C_2^0 X\frac{\partial X}{\partial t} + \left(\frac{Z_2}{R_p} + \frac{\rho_t}{\rho_t^{\text{equiv.}}}\rho_l Z_2^{-1}\right)X - \rho_t Z_2^{-1}l^2\frac{\partial^2 X}{\partial x^2} + 2l\frac{\partial X}{\partial x}$$
$$+\frac{l^3}{3}\frac{\partial^3 X}{\partial x^3} = \frac{R_t}{R_p}C_1^0 Z_1^{1/2}Z_2^{1/2}\left(\frac{\partial U}{\partial t} - 2b_1 Z_1^{1/2}U\frac{\partial U}{\partial t}\right) + \frac{Z_1^{1/2}Z_2^{1/2}}{R_p}\left(1 + \frac{\rho_t R_l}{Z_1 Z_2}\right)U \quad \text{(S.28)}$$

Multiplying equation S.27 by $\frac{\sqrt{C_2^0 Z_2}}{\sqrt{C_1^0 Z_1}}$ and equation S.28 by $\frac{\sqrt{C_1^0 Z_1}}{\sqrt{C_2^0 Z_2}}$ , we obtain

$$\sqrt{C_1^0 Z_1 C_2^0 Z_2}\frac{\partial U}{\partial t} - 2b_1 \frac{\sqrt{C_2^0 Z_2}}{\sqrt{C_1^0 Z_1}} Z_1^{3/2} C_1^0 U \frac{\partial U}{\partial t} + \left(\frac{Z_1}{R_p}\frac{R_t^{\text{equiv.}}}{R_t} + R_l Z_1^{-1}\right)\frac{\sqrt{C_2^0 Z_2}}{\sqrt{C_1^0 Z_1}} U$$

$$- R_t^{\text{equiv.}} \frac{\sqrt{C_2^0 Z_2}}{\sqrt{C_1^0 Z_1}} Z_1^{-1} l^2 \frac{\partial^2 U}{\partial x^2} + 2\frac{R_t^{\text{equiv.}}}{R_t}\frac{\sqrt{C_2^0 Z_2}}{\sqrt{C_1^0 Z_1}} l \frac{\partial U}{\partial x} + \frac{l^3}{3}\frac{R_t^{\text{equiv.}}}{R_t}\frac{\sqrt{C_2^0 Z_2}}{\sqrt{C_1^0 Z_1}}\frac{\partial^3 U}{\partial x^3}$$

$$= \frac{\rho_t}{R_p}\frac{R_t^{\text{equiv.}}}{R_t}\frac{C_2^0 Z_2^{1/2}}{C_1^0 Z_1^{1/2}}\sqrt{C_2^0 Z_2 C_1^0 Z_1}\left(\frac{\partial X}{\partial t} + 2b_2 Z_2^{1/2} X \frac{\partial X}{\partial t}\right)$$

$$+ \frac{R_t^{\text{equiv.}}}{R_t}\frac{Z_1^{1/2} Z_2^{1/2}}{R_p}\frac{\sqrt{C_2^0 Z_2}}{\sqrt{C_1^0 Z_1}}\left(1 + \frac{R_t \rho_l}{Z_1 Z_2}\right) X \quad \text{(S.29)}$$

$$\sqrt{C_1^0 Z_1 C_2^0 Z_2}\frac{\partial X}{\partial t} + 2b_2 \frac{\sqrt{C_1^0 Z_1}}{\sqrt{C_2^0 Z_2}} Z_2^{3/2} C_2^0 X \frac{\partial X}{\partial t} + \left(\frac{Z_2}{R_p}\frac{\rho_t^{\text{equiv.}}}{\rho_t} + \rho_l Z_2^{-1}\right)\frac{\sqrt{C_1^0 Z_1}}{\sqrt{C_2^0 Z_2}} X$$

$$- \rho_t^{\text{equiv.}} \frac{\sqrt{C_1^0 Z_1}}{\sqrt{C_2^0 Z_2}} Z_2^{-1} l^2 \frac{\partial^2 X}{\partial x^2} + 2\frac{\rho_t^{\text{equiv.}}}{\rho_t}\frac{\sqrt{C_1^0 Z_1}}{\sqrt{C_2^0 Z_2}} l \frac{\partial X}{\partial x} + \frac{l^3}{3}\frac{\rho_t^{\text{equiv.}}}{\rho_t}\frac{\sqrt{C_1^0 Z_1}}{\sqrt{C_2^0 Z_2}}\frac{\partial^3 X}{\partial x^3}$$

$$= \frac{R_t}{R_p}\frac{\rho_t^{\text{equiv.}}}{\rho_t}\frac{C_1^0 Z_1^{1/2}}{C_2^0 Z_2^{1/2}}\sqrt{C_1^0 Z_1 C_2^0 Z_2}\left(\frac{\partial U}{\partial t} - 2b_1 Z_1^{1/2} U \frac{\partial U}{\partial t}\right)$$

$$+ \frac{Z_1^{1/2} Z_2^{1/2}}{R_p}\frac{\rho_t^{\text{equiv.}}}{\rho_t}\frac{\sqrt{C_1^0 Z_1}}{\sqrt{C_2^0 Z_2}}\left(1 + \frac{\rho_t R_l}{Z_1 Z_2}\right) U \quad \text{(S.30)}$$

Applying the transformations

$$\xi = \frac{x}{\beta} - \frac{t}{\alpha}, \qquad \tau = \frac{t}{24\alpha}, \quad \alpha = \sqrt{C_1^0 Z_1 C_2^0 Z_2} > 0, \quad \beta = 2l$$

$$\frac{\partial}{\partial x} \equiv \frac{1}{\beta}\frac{\partial}{\partial \xi}$$

$$\frac{\partial^2}{\partial x^2} \equiv \frac{1}{\beta^2}\frac{\partial^2}{\partial \xi^2}, \qquad \frac{\partial^3}{\partial x^3} \equiv \frac{1}{\beta^3}\frac{\partial^3}{\partial \xi^3}$$

$$\frac{\partial}{\partial t} \equiv \frac{1}{\alpha}\left(\frac{1}{24}\frac{\partial}{\partial \tau} - \frac{\partial}{\partial \xi}\right)$$

and keeping the leading order terms in the parameter $\beta$, we have

$$\frac{\gamma_1}{24}\frac{\partial U}{\partial \tau}+6U\frac{\partial U}{\partial \xi}+\frac{R_t^{\text{equiv.}}}{R_t}\frac{\sqrt{C_2^0 Z_2}}{\sqrt{C_1^0 Z_1}}\frac{\gamma_1}{24}\frac{\partial^3 U}{\partial \xi^3}+\left(\frac{Z_1}{R_p}\frac{R_t^{\text{equiv.}}}{R_t}+R_l Z_1^{-1}\right)\frac{\sqrt{C_2^0 Z_2}}{\sqrt{C_1^0 Z_1}}\gamma_1 U$$
$$-R_t^{\text{equiv.}}\frac{\sqrt{C_2^0 Z_2}}{\sqrt{C_1^0 Z_1}}Z_1^{-1}\frac{\gamma_1}{4}\frac{\partial^2 U}{\partial \xi^2}+\left(\frac{R_t^{\text{equiv.}}}{R_t}\frac{\sqrt{C_2^0 Z_2}}{\sqrt{C_1^0 Z_1}}-1\right)\gamma_1\frac{\partial U}{\partial \xi}=-\frac{\rho_t}{R_p}\frac{R_t^{\text{equiv.}}}{R_t}\frac{C_2^0 Z_2^{1/2}}{C_1^0 Z_1^{1/2}}\gamma_1\frac{\partial X}{\partial \xi}$$
$$+\frac{R_t^{\text{equiv.}}}{R_t}\frac{Z_1^{1/2}Z_2^{1/2}}{R_p}\frac{\sqrt{C_2^0 Z_2}}{\sqrt{C_1^0 Z_1}}\left(1+\frac{R_t\rho_l}{Z_1 Z_2}\right)\gamma_1 X \quad \text{(S.31)}$$

$$\frac{\gamma_2}{24}\frac{\partial X}{\partial \tau}-6X\frac{\partial X}{\partial \xi}+\frac{\rho_t^{\text{equiv.}}}{\rho_t}\frac{\sqrt{C_1^0 Z_1}}{\sqrt{C_2^0 Z_2}}\frac{\gamma_2}{24}\frac{\partial^3 X}{\partial \xi^3}+\left(\frac{Z_2}{R_p}\frac{\rho_t^{\text{equiv.}}}{\rho_t}+\rho_l Z_2^{-1}\right)\frac{\sqrt{C_1^0 Z_1}}{\sqrt{C_2^0 Z_2}}\gamma_2 X$$
$$-\rho_t^{\text{equiv.}}\frac{\sqrt{C_1^0 Z_1}}{\sqrt{C_2^0 Z_2}}Z_2^{-1}\frac{\gamma_2}{4}\frac{\partial^2 X}{\partial \xi^2}+\left(\frac{\rho_t^{\text{equiv.}}}{\rho_t}\frac{\sqrt{C_1^0 Z_1}}{\sqrt{C_2^0 Z_2}}-1\right)\gamma_2\frac{\partial X}{\partial \xi}=-\frac{R_t}{R_p}\frac{\rho_t^{\text{equiv.}}}{\rho_t}\frac{C_1^0 Z_1^{1/2}}{C_2^0 Z_2^{1/2}}\gamma_2\frac{\partial U}{\partial \xi}$$
$$+\frac{Z_1^{1/2}Z_2^{1/2}}{R_p}\frac{\rho_t^{\text{equiv.}}}{\rho_t}\frac{\sqrt{C_1^0 Z_1}}{\sqrt{C_2^0 Z_2}}\left(1+\frac{\rho_t R_l}{Z_1 Z_2}\right)\gamma_2 U \quad \text{(S.32)}$$

where,

$$\gamma_1=\frac{3}{b_1 Z_1^{1/2}}$$

$$\gamma_2=\frac{3}{b_2 Z_2^{1/2}}$$

Rescaling the voltages $U=-\frac{\gamma_1}{24}\Psi$ and $X=\frac{\gamma_2}{24}\Phi$, equations S.31 and S.32 become equations 9 and 10 in the main text, where

$$N_1=24\left(\frac{Z_1}{R_p}\frac{R_t^{\text{equiv.}}}{R_t}+R_l Z_1^{-1}\right)\frac{\sqrt{C_2^0 Z_2}}{\sqrt{C_1^0 Z_1}} \quad \text{(S.33)}$$

$$N_2=24\frac{R_t^{\text{equiv.}}}{R_t}\frac{Z_1^{1/2}Z_2^{1/2}}{R_p}\frac{\sqrt{C_2^0 Z_2}}{\sqrt{C_1^0 Z_1}}\left(1+\frac{R_t\rho_l}{Z_1 Z_2}\right)\frac{\gamma_2}{\gamma_1} \quad \text{(S.34)}$$

$$N_3 = 24\left(\frac{Z_2}{R_p}\frac{\rho_t^{\text{equiv.}}}{\rho_t} + \rho_l Z_2^{-1}\right)\frac{\sqrt{C_1^0 Z_1}}{\sqrt{C_2^0 Z_2}} \tag{S.35}$$

$$N_4 = 24\frac{\rho_t^{\text{equiv.}}}{\rho_t}\frac{Z_1^{1/2}Z_2^{1/2}}{R_p}\frac{\sqrt{C_1^0 Z_1}}{\sqrt{C_2^0 Z_2}}\left(1+\frac{\rho_t R_l}{Z_1 Z_2}\right)\frac{\gamma_1}{\gamma_2} \tag{S.36}$$

$$\nu_1 = 6R_t^{\text{equiv.}}\frac{\sqrt{C_2^0 Z_2}}{\sqrt{C_1^0 Z_1}}Z_1^{-1} \tag{S.37}$$

$$\nu_2 = 6\rho_t^{\text{equiv.}}\frac{\sqrt{C_1^0 Z_1}}{\sqrt{C_2^0 Z_2}}Z_2^{-1} \tag{S.38}$$

$$M_1 = 24\left(\frac{R_t^{\text{equiv.}}}{R_t}\frac{\sqrt{C_2^0 Z_2}}{\sqrt{C_1^0 Z_1}} - 1\right) \tag{S.39}$$

$$M_2 = 24\frac{\rho_t}{R_p}\frac{R_t^{\text{equiv.}}}{R_t}\frac{C_2^0 Z_2^{1/2}}{C_1^0 Z_1^{1/2}}\frac{\gamma_2}{\gamma_1} \tag{S.40}$$

$$M_3 = 24\left(\frac{\rho_t^{\text{equiv.}}}{\rho_t}\frac{\sqrt{C_1^0 Z_1}}{\sqrt{C_2^0 Z_2}} - 1\right) \tag{S.41}$$

$$M_4 = 24\frac{R_t}{R_p}\frac{\rho_t^{\text{equiv.}}}{\rho_t}\frac{C_1^0 Z_1^{1/2}}{C_2^0 Z_2^{1/2}}\frac{\gamma_1}{\gamma_2} \tag{S.42}$$

$$F_1 = \frac{R_t^{\text{equiv.}}}{R_t}\frac{\sqrt{C_2^0 Z_2}}{\sqrt{C_1^0 Z_1}} - 1 \tag{S.43}$$

$$F_2 = \frac{\rho_t^{\text{equiv.}}}{\rho_t}\frac{\sqrt{C_1^0 Z_1}}{\sqrt{C_2^0 Z_2}} - 1 \tag{S.44}$$

### Coupled ODE system

#### Derivation of the equation for $\frac{d\kappa_1}{d\tau}$ and $\frac{d\kappa_2}{d\tau}$

We substitute equations 13 and 14 into equation 11 in the main text to get

$$P_1\text{sech}^2 y_1 = -4M_1\kappa_1^3\text{sech}^4 y_1\tanh y_1 + 4M_2\kappa_2^3\text{sech}^2 y_1\text{sech}^2 y_2\tanh y_2 + 2N_1\kappa_1^2\text{sech}^4 y_1 + 2N_2\kappa_2^2\text{sech}^2 y_1\text{sech}^2 y_2 + 4\kappa_1^4\nu_1\text{sech}^6 y_1 - 8\kappa_1^4\nu_1\text{sech}^4 y_1\tanh^2 y_1 + 32\kappa_1^5 F_1\text{sech}^6 y_1\text{tanh} y_1 - 16\kappa_1^5 F_1\text{sech}^4 y_1\text{tanh}^3 y_1 \quad \text{(S.45)}$$

Additionally, we have

$$\int_{-\infty}^{+\infty} P_1\text{sech}^2 y_1 dy_1 = \kappa_1^2[\frac{32}{15}\kappa_1^2\nu_1 + \frac{8}{3}N_1] + \int_{-\infty}^{+\infty}\text{sech}^2 y_1\text{sech}^2 y_2[2N_2\kappa_2^2 + 4M_2\kappa_2^3\tanh y_2]dy_1 \quad \text{(S.46)}$$

Substitution of equation S.46 into equation 15 in the main text, yields

$$\frac{d\kappa_1}{d\tau} = -\frac{\kappa_1}{4}[\frac{32}{15}\kappa_1^2\nu_1 + \frac{8}{3}N_1] - \frac{\kappa_2^3}{\kappa_1}\int_{-\infty}^{\infty}\text{sech}^2 y_1\text{sech}^2 y_2[\frac{N_2}{2\kappa_2} + M_2\tanh y_2]dy_1 \quad \text{(S.47)}$$

where

$$y_2 = \frac{\kappa_2}{\kappa_1}y_1 + \kappa_2\Delta\theta$$

Following a similar procedure, we have

$$P_2\text{sech}^2 y_2 = -4M_3\kappa_2^3\text{sech}^4 y_2\tanh y_2 + 4M_4\kappa_1^3\text{sech}^2 y_1\text{sech}^2 y_2\tanh y_1 + 2N_3\kappa_2^2\text{sech}^4 y_2 + 2N_4\kappa_1^2\text{sech}^2 y_1\text{sech}^2 y_2 + 4\kappa_2^4\nu_2\text{sech}^6 y_2 - 8\kappa_2^4\nu_2\text{sech}^4 y_2\tanh^2 y_2 + 32\kappa_2^5 F_2\text{sech}^6 y_2\text{tanh} y_2 - 16\kappa_2^5 F_2\text{sech}^4 y_2\text{tanh}^3 y_2 \quad \text{(S.48)}$$

$$\int_{-\infty}^{+\infty} P_2\text{sech}^2 y_2 dy_2 = \kappa_2^2[\frac{32}{15}\kappa_2^2\nu_2 + \frac{8}{3}N_3] + \int_{-\infty}^{+\infty}\text{sech}^2 y_1\text{sech}^2 y_2[2N_4\kappa_1^2 + 4M_4\kappa_1^3\tanh y_1]dy_2 \quad \text{(S.49)}$$

and

$$\frac{d\kappa_2}{d\tau} = -\frac{\kappa_2}{4}[\frac{32}{15}\kappa_2^2\nu_2 + \frac{8}{3}N_3] - \frac{\kappa_1^3}{\kappa_2}\int_{-\infty}^{\infty}\text{sech}^2 y_1\text{sech}^2 y_2[\frac{N_4}{2\kappa_1} + M_4\tanh y_1]dy_2 \quad \text{(S.50)}$$

where,

$$y_1 = \frac{\kappa_1}{\kappa_2}y_2 - \kappa_1\Delta\theta$$

### Derivation of the equations for $\frac{d\theta_1}{d\tau}$ and $\frac{d\theta_2}{d\tau}$

We substitute of the following identity

$$P_1(y_1+\frac{1}{2}\sinh 2y_1)\text{sech}^2 y_1 = P_1 y_1 \text{sech}^2 y_1 + P_1 \tanh y_1 \tag{S.51}$$

into equation 16 in the main text. Further substitution of equations 13 and 14 into the previous equation yields

$$\begin{aligned} P_1 y_1 \text{sech}^2 y_1 = & -4M_1\kappa_1^3 y_1 \text{sech}^4 y_1 \tanh y_1 + 4M_2\kappa_2^3 y_1 \text{sech}^2 y_1 \text{sech}^2 y_2 \tanh y_2 + 2N_1\kappa_1^2 y_1 \text{sech}^4 y_1 \\ & + 2N_2\kappa_2^2 y_1 \text{sech}^2 y_1 \text{sech}^2 y_2 + 4\kappa_1^4 \nu_1 y_1 \text{sech}^6 y_1 - 8\kappa_1^4 \nu_1 y_1 \text{sech}^4 y_1 \tanh^2 y_1 + 32\kappa_1^5 F_1 y_1 \text{sech}^6 y_1 \tanh y_1 \\ & - 16\kappa_1^5 F_1 y_1 \text{sech}^4 y_1 \tanh^3 y_1 \end{aligned} \tag{S.52}$$

and

$$\begin{aligned} P_1 \tanh y_1 = & -4M_1\kappa_1^3 \text{sech}^2 y_1 \tanh^2 y_1 + 4M_2\kappa_2^3 \text{sech}^2 y_2 \tanh y_2 \tanh y_1 + 2N_1\kappa_1^2 \text{sech}^2 y_1 \tanh y_1 \\ & + 2N_2\kappa_2^2 \text{sech}^2 y_2 \tanh y_1 + 4\kappa_1^4 \nu_1 \text{sech}^4 y_1 \tanh y_1 - 8\kappa_1^4 \nu_1 \text{sech}^2 y_1 \tanh^3 y_1 + 32\kappa_1^5 F_1 \text{sech}^4 y_1 \tanh^2 y_1 \\ & - 16\kappa_1^5 F_1 \text{sech}^2 y_1 \tanh^4 y_1 \end{aligned} \tag{S.53}$$

Additionally, we have

$$\int_{-\infty}^{+\infty} P_1 y_1 \text{sech}^2 y_1 dy_1 = -\frac{4}{3}M_1\kappa_1^3 + \kappa_2^3 \int_{-\infty}^{+\infty} y_1 \text{sech}^2 y_1 \text{sech}^2 y_2 [\frac{2N_2}{\kappa_2} + 4M_2 \tanh y_2] dy_1 + \frac{16}{5}\kappa_1^5 F_1 \tag{S.54}$$

and

$$\int_{-\infty}^{+\infty} P_1 \tanh y_1 dy_1 = -\frac{8}{3}M_1\kappa_1^3 + \frac{32}{15}\kappa_1^5 F_1 + \kappa_2^3 \int_{-\infty}^{+\infty} \text{sech}^2 y_2 \tanh y_1 [\frac{2N_2}{\kappa_2} + 4M_2 \tanh y_2] dy_1 \tag{S.55}$$

As a result, we obtain

$$\begin{aligned} \int_{-\infty}^{+\infty} P_1(y_1+\frac{1}{2}\sinh 2y_1)\text{sech}^2 y_1 dy_1 = & -4M_1\kappa_1^3 + \frac{16}{3}\kappa_1^5 F_1 \\ & + \kappa_2^3 \int_{-\infty}^{+\infty} [y_1 \text{sech}^2 y_1 + \tanh y_1] \text{sech}^2 y_2 [\frac{2N_2}{\kappa_2} + 4M_2 \tanh y_2] dy_1 \end{aligned} \tag{S.56}$$

Substitution of the previous equation into equation 16 in the main text, we get

$$\frac{d\theta_1}{d\tau} = 4\kappa_1^2 + M_1 - \frac{4}{3}\kappa_1^2 F_1 - \frac{\kappa_2^3}{4\kappa_1^3}\int_{-\infty}^{+\infty}[y_1 \mathrm{sech}^2 y_1 + \tanh y_1]\mathrm{sech}^2 y_2 [\frac{2N_2}{\kappa_2} + 4M_2 \tanh y_2] dy_1 \qquad \text{(S.57)}$$

Following a similar procedure, we get,

$$\frac{d\theta_2}{d\tau} = 4\kappa_2^2 + M_3 - \frac{4}{3}\kappa_2^2 F_2 - \frac{\kappa_1^3}{4\kappa_2^3}\int_{-\infty}^{+\infty}[y_2 \mathrm{sech}^2 y_2 + \tanh y_2]\mathrm{sech}^2 y_1 [\frac{2N_4}{\kappa_1} + 4M_4 \tanh y_1] dy_2 \qquad \text{(S.58)}$$

### Numerical solution of the coupled ODE system

We used Mathematica 14.0 and the default settings for the numerical solver NDSolve to solve equations S.47, S.50, S.57, and S.58 numerically. The initial conditions are given by the expressions $\kappa_1(0) = \sqrt{\frac{12V_{\mathrm{inp}}}{\gamma_1\sqrt{Z_1}}}$, $\kappa_2(0) = \sqrt{\frac{12W_{\mathrm{inp}}}{\gamma_2\sqrt{Z_2}}}$, and we considered the initial soliton phase conditions used in ref. [1], namely $\theta_1(0) = 0.25$, and $\theta_2(0) = 0.20$. We also set $\kappa = 0.5\,[\kappa_1(0) + \kappa_2(0)]$ and solved the equations system for input voltages $V_{\mathrm{inp}} = W_{\mathrm{inp}}$ between 10 mV and 100 mV. In the automatic method setting, NDSolve chooses the best-suited method for the equations, which can be an explicit method (e.g., "Adams") or an implicit method (e.g., "BDF" [backward differentiation formulas]). However, both methods produced the same result for our equations system. Since we did not specify the step size, NDSolve automatically adjusted the step size as needed. The WorkingPrecision was set to the MachinePrecision since we used machine-precision numbers (double-precision floating-point numbers: 16 decimal digits). NDSolve typically uses error tolerance between $10^{-6}$ and $10^{-7}$ . The default interpolation order is usually three, and the AccuracyGoal is infinity.

### Oscillation frequency calculation

To estimate the soliton oscillation frequency, we analyzed the time-dependent voltage signal by detecting its local maxima over the dimensionless time variable $\tau$. Specifically, we evaluated the soliton voltage at a fixed spatial location and identified successive peaks in the signal using a peak detection algorithm (see Fig. 4 in the main text). The average time interval between peaks, $\Delta\tau$, was computed and used to determine the oscillation period. Since the relation between dimensionless time and physical time is given by $\tau = t/(24\alpha)$, the physical period is $T = 24\alpha\Delta\tau$, and the corresponding frequency is $f = 1/(24\alpha\Delta\tau)$. Using the model parameter $\alpha = 4.265 \times 10^{-5}s$, we calculated the soliton oscillation frequency to be approximately 39 Hz. This value is in good agreement with experimental measurements obtained from voltage-clamp recordings (see Fig. 3 in the main text).

### Approximate analytical solution of the coupled ODE system

We have numerically solved the coupled ODE system–equations for multiple values of the involved parameters and concluded that the parameters $M_1 = M_3 = N_2 = N_4 = F_1 = F_2 = 0$ change the soliton amplitudes but they weakly affect their oscillation frequency. Thus, we neglected these parameters and linearized equations S.47, S.50, S.57, and S.58 for small $\Delta\theta$ to obtain an approximate analytical expression for the solitons oscillation frequency. In doing so, the coupled ODE system–equations become

$$\frac{\partial \lambda_1}{\partial \tau} = -\frac{8}{15} M_2 \kappa^3 \Delta\theta - \frac{2}{3} N_1 (\kappa + \lambda_1) \tag{S.59}$$

$$\frac{\partial \lambda_2}{\partial \tau} = \frac{8}{15} M_4 \kappa^3 \Delta\theta - \frac{2}{3} N_3 (\kappa + \lambda_2) \tag{S.60}$$

$$\frac{\partial \theta_1}{\partial \tau} = -M_2 + 4\kappa^2 + 8\kappa\lambda_1 + \frac{2M_2 \Delta\lambda \left(30 + \pi^2\right)}{45\kappa} \tag{S.61}$$

$$\frac{\partial \theta_2}{\partial \tau} = -M_4 + 4\kappa^2 + 8\kappa\lambda_2 - \frac{2M_4 \Delta\lambda \left(30 + \pi^2\right)}{45\kappa} \tag{S.62}$$

where $\kappa_1 = \kappa + \lambda_1$, $\kappa_2 = \kappa + \lambda_2$, and $\Delta\lambda = \lambda_1 - \lambda_2$. From equations S.59 and S.60,

$$\frac{\partial \Delta\lambda}{\partial \tau} = -\frac{8}{15} (M_2 + M_4) \kappa^3 \Delta\theta - \frac{2}{3} N \Delta\lambda \tag{S.63}$$

In which we set $N_1 = N_3 = N$.

Similarly, from equations S.61 and S.62, we can write

$$\frac{\partial \Delta\theta}{\partial \tau} = -(M_2 - M_4) + \left[8\kappa + \frac{2}{45\kappa}\left(30 + \pi^2\right)(M_2 + M_4)\right] \Delta\lambda \tag{S.64}$$

From equation S.64, we get

$$\frac{\partial^2 \Delta\theta}{\partial \tau^2} = A \frac{\partial \Delta\lambda}{\partial \tau} \tag{S.65}$$

Where, $A = 8\kappa + \frac{2}{45\kappa}\left(30 + \pi^2\right)(M_2 + M_4)$.

Substitution of equation S.63 into equation S.65 yields

$$\frac{\partial^2 \Delta\theta}{\partial \tau^2} = -\frac{8A}{15} (M_2 + M_4) \kappa^3 \Delta\theta - \frac{2}{3} N (A \Delta\lambda) \tag{S.66}$$

Substitution of equation S.64 into equation S.66 provides the equation 17 in the main text.

Additionally, we get from equation S.63

$$\frac{\partial^2 \Delta\lambda}{\partial \tau^2} = -\frac{8}{15} (M_2 + M_4) \kappa^3 \frac{\partial \Delta\theta}{\partial \tau} - \frac{2}{3} N \frac{\partial \Delta\lambda}{\partial \tau} \tag{S.67}$$

Substitution of equation S.64 into equation S.67 generates the equation 20 in the main text.

Once we have the solutions for $\Delta\theta$ and $\Delta\lambda$, we can write

$$\frac{\partial \lambda_1}{\partial \tau} = -\frac{8}{15}M_2\kappa^3\Delta\theta - \frac{2}{3}N_1\kappa - \frac{2}{3}N_1\lambda_1 \tag{S.68}$$

Defining $L = \frac{2}{3}N_1$, and $-\frac{8}{15}M_2\kappa^3\Delta\theta = f(\tau)$, equation S.68 becomes

$$\frac{\partial \lambda_1}{\partial \tau} = -L\kappa - L\lambda_1 + f(\tau) \tag{S.69}$$

Setting $\lambda_1 = \kappa_1 - \kappa$, we have

$$\frac{\partial k_1}{\partial \tau} = -L\kappa_1 + f(\tau) \tag{S.70}$$

and using $\kappa_1 = p_1 e^{-L\tau}$, $\frac{\partial \kappa_1}{\partial \tau} = \frac{\partial p_1}{\partial \tau}e^{-L\tau} - L\kappa_1$, we get

$$\frac{\partial p_1}{\partial \tau} = f(\tau)e^{L\tau} \tag{S.71}$$

Thus, we obtain the solution

$$p_1 = \int d\tau f(\tau)e^{L\tau} + C \tag{S.72}$$

and

$$\lambda_1 = -\kappa + p_1 e^{-L\tau}$$

$$\lambda_1 = -\kappa + \left[\int d\tau f(\tau)e^{L\tau} + C\right]e^{-L\tau} \tag{S.73}$$

Since at $\tau = 0$, $\lambda_0 = -\kappa + p_1(0) + C$, then $C = \lambda_0 + \kappa - p_1(0)$ and

$$\lambda_1 = -\kappa + \left[\int d\tau f(\tau)e^{L\tau} + \lambda_0 + \kappa - p_1(0)\right]e^{-L\tau} \tag{S.74}$$
